# Mechano-calcium and mechano-electric feedbacks in the human cardiomyocyte analyzed in a mathematical model

**DOI:** 10.1101/855890

**Authors:** Nathalie A. Balakina-Vikulova, Alexander Panfilov, Olga Solovyova, Leonid B. Katsnelson

**Affiliations:** Institute of Immunology and Physiology of the Ural Branch of the Russian Academy of Sciences, Ekaterinburg, Russia; Ural Federal University, Ekaterinburg, Russia; Ghent University, Ghent, Belgium

**Keywords:** Human myocardium, Electromechanical coupling, Mechano-electric feedback

## Abstract

Experiments on animal hearts (rat, rabbit, guinea pig, etc.) have demonstrated that mechano-calcium feedback (MCF) and mechano-electric feedback (MEF) are very important for myocardial self-regulation because they adjust the cardiomyocyte contractile function to various mechanical loads and to mechanical interactions between heterogeneous myocardial segments in the ventricle walls. In *in vitro* experiments on these animals, MCF and MEF manifested themselves in several basic classical phenomena (e.g. load dependence, length dependence of isometric twitches, etc.), and in the respective responses of calcium transients and action potentials. However, it is extremely difficult to study simultaneously the electrical, calcium, and mechanical activities of the human heart muscle *in vitro*. Mathematical modeling is a useful tool for exploring these phenomena. We have developed a novel model to describe electromechanical coupling and mechano-electric feedbacks in the human cardiomyocyte. It combines the ‘ten Tusscher – Panfilov’ electrophysiological model of the human cardiomyocyte with our module of myocardium mechanical activity taken from the ‘Ekaterinburg – Oxford’ model and adjusted to human data. Using it, we simulated isometric and afterloaded twitches and effects of MCF and MEF on excitation-contraction coupling. MCF and MEF were found to affect significantly the duration of the calcium transient and action potential in the human cardiomyocyte model in response to both smaller afterloads as compared to bigger ones and various mechanical interventions applied during isometric and afterloaded twitches.

## 1 Introduction

Although the main function of electrical excitation in the heart is to trigger mechanical contraction, the latter influences in turn electrical wave propagation via processes called mechano-calcium feedbacks (MCF) and mechano-electric feedbacks (MEF) [1–3]. Physiologically, mechanical conditions ensure the adaptation of the normal myocardium to varying external and internal mechanical conditions of contraction, including global and local mechanical loading and length redistribution between interacting heterogeneous cardiomyocytes in the heart chamber walls [4]. Interacting myocardial cells differ in the duration of twitches and in the ability to develop mechanical tension, and are excited with a delay relative to each other, and these differences create dynamically changing mechanical loading conditions for them. The result is a complex continuous chain of feedbacks (MEF and MCF) between the contraction of the entire ensemble of cardiomyocytes in the wall and activation of each of them. These feedbacks dynamically modulate the subsequent course of the twitch in each cardiomyocyte. Previously, we studied these interaction effects experimentally on the simplest biological models: heterogeneous muscle duplexes [5–8] and mathematical models for such duplexes, and one-dimensional models of heterogeneous cardiac tissue [5, 6, 8–10].

On the other hand, the pathological manifestations of these feedbacks can be quite dramatic: for example, they can cause heart rhythm disturbances and even lead to sudden cardiac death [1]. Experimental data on electromechanical activity are available mostly for animal hearts, while data for the human cardiomyocyte are very limited. Neither have the mechanisms behind MCF and MEF been fully understood.

In particular, several intracellular mechanisms have been shown to participate in the cardiac MCF. The mechano-dependence of the sarcoplasmic reticulum Ca^2+^ spark rate may contribute to the MCF via calcium-induced calcium release [11]. A key MCF circuit is associated with the load and/or length dependent kinetics of calcium-troponin C (TnC) complexes regulating Ca^2+^ activation of cardiomyocyte contractions [12, 13].

MCF contributes to both fast and slow force responses of the heart muscle to stretches and other types of deformation. Moreover, MCF can trigger MEF since the mechano-dependent modulation of Ca^2+^ transient shape and duration during a mechanical twitch can influence the shape and duration of the action potential via Ca^2+^ currents including the Na^+^-Ca^2+^ exchange one.

Another mechanism of the MEF is related to the activity of mechano-sensitive ion channels in the cardiomyocyte membrane [14]. While the contribution of these channels to MEF is generally recognized, many details remain poorly understood, including the characteristics of their conductivity response to the stretch, their localization in the cell membrane, and ionic specificity. There are reasons to believe that the contribution of stretch-activated ion channels to myocardium mechanical activity is more likely to manifest itself in slow force response effects rather than fast ones (see Discussion). In this work, we focus mainly on the fast force response phenomena listed below.

In experiments on isolated cardiac muscle preparations (trabeculae and papillary muscles) from animal hearts (rat, rabbit, guinea pig, etc.), MCF and MEF were revealed as the following basic classical phenomena: length dependence of isometric twitches, load dependence of afterloaded contractions, muscle inactivation due to short-time deformations during isometric twitches (see [15] for a review), and as respective responses of calcium transients and action potentials to mechanical interventions. Scarce data of this kind are available for human cardiac preparations whereas the contraction-excitation feedback in humans, mechano-dependent arrhythmias included, has been supported by abundant clinical evidence [1, 16, 17].

The prime objective of this paper is to suggest a way to study and predict possible MCF and MEF manifestations in the human cardiac muscle by means of modeling. To this end, we have developed a novel mathematical electromechanical model by integrating limited experimental data on excitation-contraction and mechano-electrical coupling in healthy and pathological human myocardium. The electromechanical model of the human cardiomyocyte is an important result in itself as it can be used to examine electromechanical effects in the myocardial tissue and in the whole human heart for diagnosis and effective treatment of cardiac diseases.

Several models have been developed for describing the electrophysiological function of human cardiomyocytes [18–21]. However, few of these models are combined with cardiac mechanics [22–25]. The question of the presence of MEF in human cardiomyocytes was raised in the recent work [24], where two human electrophysiological models [19, 21] were combined with mechanical models developed by J. Rice [26] and by S. Land [27]. The former parametrically fitted his model to the human cardiomyocyte whereas Land developed a model specially to simulate contractions and passive viscoelasticity in it. Land’s model inherited MCF in electromechanical models from the Rice model as well. The authors did not discover MEF in any of the electromechanical models they considered and therefore concluded that MEF are unlikely to be present in the healthy human myocardium. This conclusion, however, is based on the specific features of MCF in Rice model [26], while it is likely that it cannot properly describe the wide range of important mechanics effects. In the ‘Discussion’, we will characterize in more detail some important distinctions between our approach and that of the Rice model that relate to the difference in the description of cardiomyocyte calcium activation mechano-sensitivity.

Experimental studies have proved that MEF does take place in the myocardium of many mammalian species, from the cellular to organ level, and manifests itself in the intact human hearts *in vivo*. We thus believe that this property should also be present in human myocardial cells and play an important physiological/pathophysiological role in the regulation of the human heart function. In this paper, we report a novel electromechanical model combining the electrophysiological model [20] and the model of cardiomyocyte mechanics [28]. Using it, we evaluate MCF and MEF manifestations in human cardiac cells.

## 2 Methods

### 2.1 TP+M model

Our combined model of excitation-contraction coupling (ECC) in the human cardiomyocyte (TP+M model) is formed of two main modules: an electrophysiological and a mechanical part coupled to each other.

The module for cellular electrophysiology is based on the 2006 version of the ‘ten Tusscher – Panfilov’ (TP) ionic model, which is described in detail elsewhere [20] and has been widely used in electrophysiological studies [20, 29–32]. The TP model describes ionic currents across the membrane and generation of the action potential (AP) (see Fig. 1 for details). In particular, this model was used as the electrical component in electromechanical models combined with different mechanical modules [22, 23, 33].

**Fig. 1.**
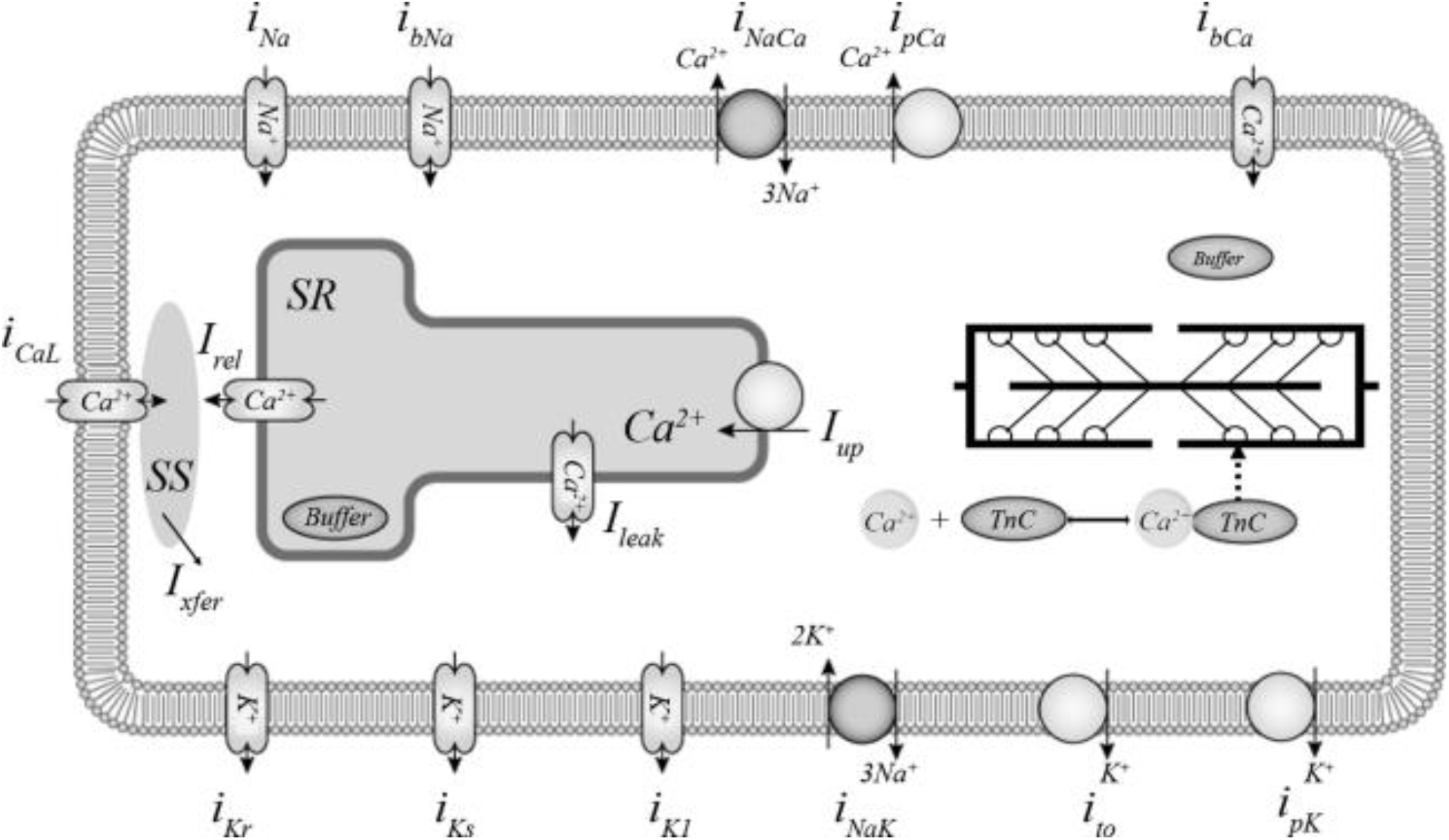
Ionic membrane currents and intracellular calcium homeostasis in the TP+M model. **Calcium currents** : *i*_*CaL*_ – L-type Ca^2+^ current; *i*_*bCa*_ – background Ca^2+^ current. **Potassium currents** : *i*_*K1*_ – inward rectifier K^+^ current; *i*_*to*_ – transient outward current; *i*_*Kr*_, *i*_*Ks*_ – rapid and slow delayed rectifier current; *i*_*pK*_ – plateau K^+^ current. **Sodium currents** : *i*_*Na*_ – fast Na^+^ current; *i*_*bNa*_ – background Na^+^ current. **Pumps and exchangers** : *i*_*pCa*_ – sarcolemmal Ca^2+^ pump current; *i*_*NaK*_ – Na^+^-K^+^ pump current; *i*_*NaCa*_ – Na^+^-Ca^2+^ exchanger (NCX) current. **Calcium translocations** : *I*_*rel*_ – Ca^2+^ release from the sarcoplasmic reticulum (SR) via ryanodine receptors to the subspace (SS); *I*_*xfer*_ – Ca^2+^ diffusion from SS to the cytoplasm; *I*_*leak*_ – a small Ca^2+^ leakage from the SR to the cytoplasm; *I*_*up*_ – Ca^2+^ pumping from the cytoplasm to the SR, where Ca^2+^ is partially buffered (*Buffer* in SR). Cytoplasmic buffering is divided into two compartments: Ca^2+^-troponin C complex formation (*Ca-TnC*) inherited from [28] and buffering by other intracellular ligands (*Buffer*). The figure is modified from the diagram in the Physiome Model Repository^4^ based on [20].

The module of mechanical activity was developed by us previously and used as a component in the ‘Ekaterinburg – Oxford’ electromechanical model [28] describing ECC in various animal cardiomyocytes (e.g. guinea pig and rabbit). It is adaptable to each particular species via parameter tuning.

The rheological scheme of this module is presented in Fig. 2. The mechanics of the passive elastic and viscous elements, as well as the mechanical activity of the contractile element are described by a system of ordinary differential and algebraic equations we published previously [28]. The only difference from that work is that the new model has two viscous elements to support the simulation of cardiac biomechanical effects observed experimentally. As we had shown even before the cited article by Sulman et al., the second viscous element (see Fig. 2) is responsible for the experimentally observed time lag between the duration of the muscle force increase phase and the duration of sarcomere shortening during isometric twitches [34]. This property turns out quite useful for fitting the time-to-peak isometric force to the range of the experimentally observed values (see below). In other words, in the article by Sulman et al. the second viscous element was excluded only for simplifying the model; however, in cases where the model should simulate quantitative species-specific features of the contractile cycle, this element cannot be neglected.

**Fig. 2.**
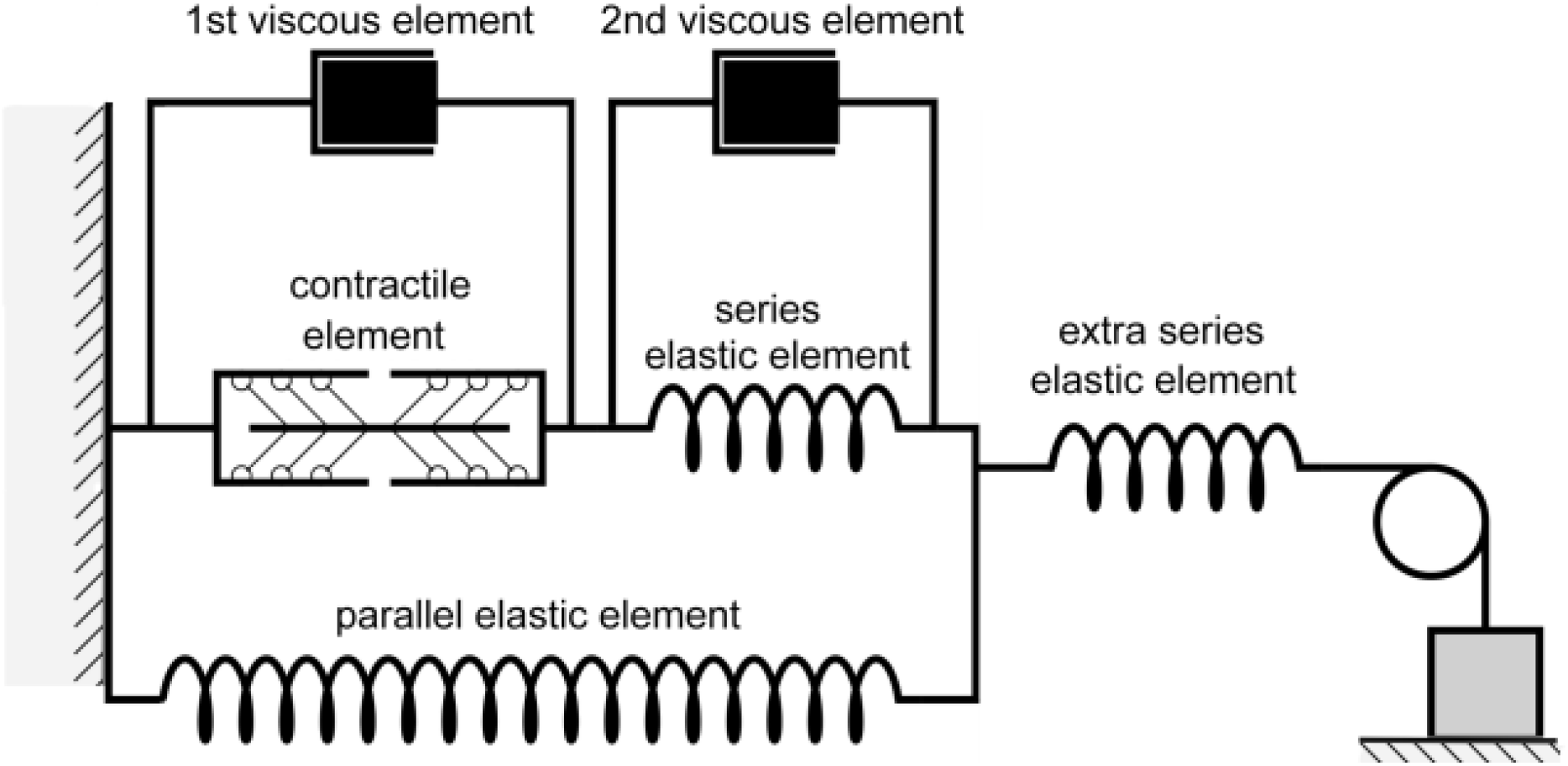
Rheological scheme of virtual cardiac sample in the TP+M model. It consists of a contractile element, which is a generator of active force, three elastic and two viscous elements. Contractile element in the model reproduces sarcomeres in the cardiomyocyte

The main component of the rheological scheme is the contractile element representing sarcomeres in the cardiomyocyte. This element is responsible for both generation of active force developed by the cardiomyocyte and its shortening due to the cross-bridges (Xb-s) formed by myosin heads attached to the actin thin filaments. Xb attachment during the contractile cycle is regulated by Ca^2+^ ions through their binding to TnC located along the thin filament. The differential equations describing Xb kinetics and Ca^2+^-troponin C complex (Ca-TnC) kinetics in the model are provided in the Supplement.

An important feature of this model is the mathematical description of the mechanisms of cooperativity [20, 28], which make the kinetics of Ca-TnC dependent on the number of attached force-generating Xb-s. In particular, the decay of the Ca-TnC complex becomes slower if a larger number of Xb-s bind to actin and/or if a larger number of other Ca-TnC complexes are formed along the thin filament near it.

The cooperative effects of an attached Xb on Ca-TnC kinetics contribute to the MCF in the ‘Ekaterinburg – Oxford’ model via the length dependence of Xb attachment/detachment in the model.

Particular mechanisms underlying the effects of length on Xb attachment have been the subject of much debate. For a long time, the prevailing concept was that of lattice spacing suggesting that a decrease in the sarcomere inter-filament distance due to cardiomyocyte lengthening facilitates actomyosin interactions and thus increases the probability of Xb attachment [28, 35, 36]. Recent experimental studies have unraveled an essential role of the giant protein titin in the length dependence of Xb dynamics [37, 38]. In the ‘Ekaterinburg – Oxford’ model, the length dependent probability of Xb binding is described phenomenologically as an overall result of all possible underlying molecular mechanisms. Thus, this formalization is invariant with respect to the subtle mechanisms discussed. Together with above-mentioned cooperativity, this length dependent probability allows the ‘Ekaterinburg – Oxford’ model to reproduce most of the MCF and MEF effects found in the heart muscle.

The cooperativity of Ca^2+^ myofilament activation in the mechanical module has allowed the ‘Ekaterinburg – Oxford’ model to reproduce a wide range of experimental manifestations of the effects of mechanical conditions of cardiac muscle contraction on the intracellular calcium kinetics (in particular, the effect of its shortening). Moreover, MCF allows it to reproduce the effects of contraction on electrical cellular activity, particularly on AP shape and duration [9, 28]. Ca^2+^-dependent modulation in the Na^+^-Ca^2+^ exchange (NCX) current of the Noble’98 ionic model [39] was found to trigger MEF in the combined ‘Ekaterinburg – Oxford’ model [9, 28].

The TP ionic model and the Ekaterinburg mechanical model are coupled to form the TP+M model via Ca^2+^ handling. However, description of such coupling required some modification of the model. In particular, the TP model does not describe Ca-TnC kinetics. It contains a mathematical description only for the generalized calcium-binding buffer in the cytosol, simulating the overall contribution of calcium-binding intracellular ligands and TnC. Moreover, a quasi-stationary algebraic equation is used in this model to describe cytosolic Ca^2+^ buffering.

However, Ca-TnC kinetics is essential for contraction initiation. It is therefore described in the Ekaterinburg mechanical model by an ordinary differential equation separately from other intracellular Ca^2+^ binding ligands. Moreover, the cooperative mechanisms of MCF linking the Ca-TnC and Xb kinetics are described by corresponding differential equations in the mechanical module of the Ekaterinburg model. These cross-links are key cardiac ECC mechanisms and, therefore, our new model should take them into account to be able to evaluate the extent of MCF and MEF manifestations in human cardiac cells.

Like the Ekaterinburg mechanical module, the combined TP+M model contains a separate differential equation for Ca-TnC kinetics. We also re-fitted the parameters of the algebraic quasi-stationary equation of the generalized calcium buffer in the model under consideration to make its sum with the calcium Ca-TnC buffer relevant for the overall generalized calcium buffer of the TP model.

In addition, we changed some parameter values in the equation for the NCX current (*i*_*NaCa*_) from the TP model to fit the combined TP+M model to experimental data. The details of this parametric tuning are described below.

All the equations and parameters of the combined TP+M model are presented in the Supplement. Here we provide only the equations of the of Ca^2+^ handling module that were modified in the TP+M model compared to the TP model.

The following equation describes time-dependent changes in Ca-TnC concentration (*[Ca-TnC]*) [28]:

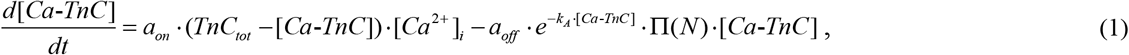

where *TnC*_*tot*_ is the total concentration of TnC in the cytosol; Π(*N*) is the cooperative dependence of Ca-TnC dissociation on Xb concentration N; *a*_*on*_, *a*_*off*_, *k*_*A*_ are model parameters (see the Supplement for details).

The cooperativity mechanisms are described in detail in our previous works [28, 40, 41].

The following equation describes time-dependent changes in free intracellular Ca^2+^ concentration (*[Ca_2+_]_i_*):

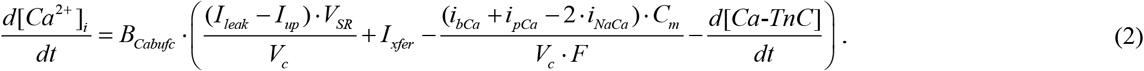

where the first term describes Ca^2+^ release (*I*_*up*_) and leakage (*I*_*leak*_) from the sarcoplasmic reticulum (SR) and Ca^2+^ SERCA uptake (*I_up_*) to the SR (*V*_*c*_ is the cytoplasmic volume, *V*_*SR*_ is the SR volume). The second term (*I*_*xfer*_) refers to Ca^2+^ diffusion leakage from the subspace into the cytoplasm. The third term describes Ca^2+^ currents through the cell membrane (*F* is the Faraday constant, *C*_*m*_ is the membrane capacitance). The descriptions of the fluxes and ion currents are inherited from the TP model. The term 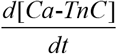 accounts for Ca^2+^ binding to TnC as provided by Eq. 1.

The factor *B*_*Cabufc*_ is derived from the quasistationary formulation for the overall concentration of Ca^2+^ bound to intracellular buffer ligands other than troponin C in the TP model, and *B*_*Cabufc*_ is non-dimensional:

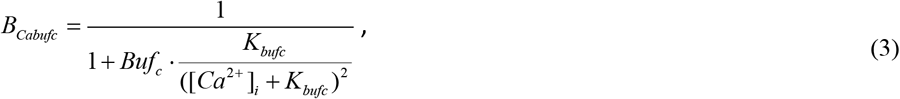

where *Buf*_*c*_ is the total concentration of all calcium-binding buffer proteins in the cytoplasm excepting TnC, *K*_*bufc*_ is the half-saturation constant for the generalized Ca^2+^ buffer.

The description of the generalized buffer represented by Eq. 3 comes from the TP model. We changed some parameters in Eq. 3 to fit the sum of the modified calcium-buffer concentration and *[Ca-TnC]* in the combined TP+M model to the overall calcium-buffer concentration in the original TP model.

The following formulation for the NCX current is also taken from the TP model:

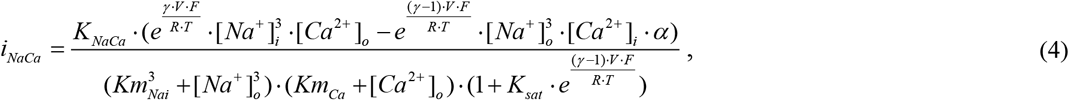

where *V* is the membrane potential, γ is the parameter for the dependence of the NCX current on the potential; *[Na*^+^]_*i*_, *[Ca*^*2*+^]_*i*_, *[Na*^+^]_*o*_, *[Ca*^*2*+^]_*o*_ are intracellular and extracellular concentrations of Na^+^ and Ca^2+^; *K*_*NaCa*_ is the maximum NCX current; *Km*_*Ca*_ is the saturation constant for Ca^2+^; *Km*_*Nai*_ is the saturation constant for Na^+^; *K*_*sat*_ is the saturation coefficient for the NCX current at very negative potentials; α is a multiplier due to which one can increase the contribution of the term responsible for the inwardly directed current; *R* is the gas constant; *F* is the Faraday constant; *T* is the temperature.

### 2.2 Simulation design and protocols

The prime objective of this paper is to analyze possible effects of MCF and MEF in the human myocardium in various modes of contraction. To this end, we carried out three types of test for a virtual preparation represented by the TP+M model: (1) various loading modes applied to the contracting virtual sample; (2) different initial lengths of the virtual sample, i.e. different preloads; and (3) quick changes in length during the contractions of the sample.

As was mentioned in the Introduction, experimental data on the mechanical activity of cellular or multicellular human myocardial preparations are very limited. The few experimental works that were performed in the 1990s studied the contraction of papillary muscles and trabeculae from human heart ventricles in isometric mode [42–57]. Only two papers reported studies of contractions under mechanical loads [47, 55]. In our simulations reported below, we compared, wherever possible, the data of our numerical experiments with the results of the studies referred to above. In the latter, however, action potentials (APs) were not recorded during muscle contractions. On the other hand, the protocols of all our numerical experiments reproduce the protocols implemented in various animals experiments used for quantifying MCF and MEF. Therefore, all phenomena we model are also compared qualitatively with data from such studies on animals, including mechanical readouts, Ca^2+^ transients in cardiomyocytes, and APs.

Simulated signals generated by the TP+M model in the isometric mode of contraction are shown in the steady-state for the pacing rate of 1 Hz. Simulated responses to mechanical interventions (e.g. change in the imposed external load (afterload), quick stretch, and quick release) are evaluated during the first twitch in which the intervention is applied according to experimental protocols [41, 55, 58, 59].

Thus, our study is focused on the analysis of the fast (immediate) responses of the human myocardium to mechanical exposures.

## 3 Results

The following circumstances are related to the specifics of human myocardial model validation:

- there is significant scatter in quantitative characteristics obtained in very few experimental studies of human human myocardial contractility;
- among these studies, none recorded mechanical, electrical, and calcium signals simultaneously from one experimental sample / group of samples.

These considerations made it impossible to develop a quantitative electromechanical model for any particular or “average” human cardiomyocyte preparation. Therefore the main validation criteria for us were:

1. the possibility to find values for the model parameters such that would ensure simulated characteristics falling within experimental data ranges where the latter were available;
2. given such parameter values, the ability to reproduce qualitatively the scarce experimental observations testifying to the influence of mechanical conditions on the contractile, electrical, and calcium activity of the human heart muscle.

Following such verification, the model could be applied to reveal and analyze MEF and MCF in other numerical experiments compared to actual ones performed on myocardial preparations from various species other than humans.

Most of the experiments on the human myocardium were conducted in the isometric mode of contractions [21, 42, 45, 47, 49, 51, 52, 55, 56]. The simulation of isometric contractions should therefore be the main platform for model validation as outlined below.

### 3.1 Simulation of steady state isometric twitches. Model validation

Using the TP+M model, we simulated isometric twitches for various diastolic lengths of the virtual preparation. These simulations qualitatively fit well with the experimentally recorded isometric twitches of isolated human trabeculae when the muscle length was reduced in a stepwise fashion [56]. These authors presented mechanical readouts (length and force) only, whereas our simulations reveal (Fig. 3) mechanical and electrical signals and calcium transients obtained at various lengths (from 80%*L*_*max*_ to 95%*L*_*max*_) under steady state conditions for the pacing rate of 1 Hz. Fig. 3b shows the curves just for extreme cases (the largest and smallest lengths).

**Fig. 3.**
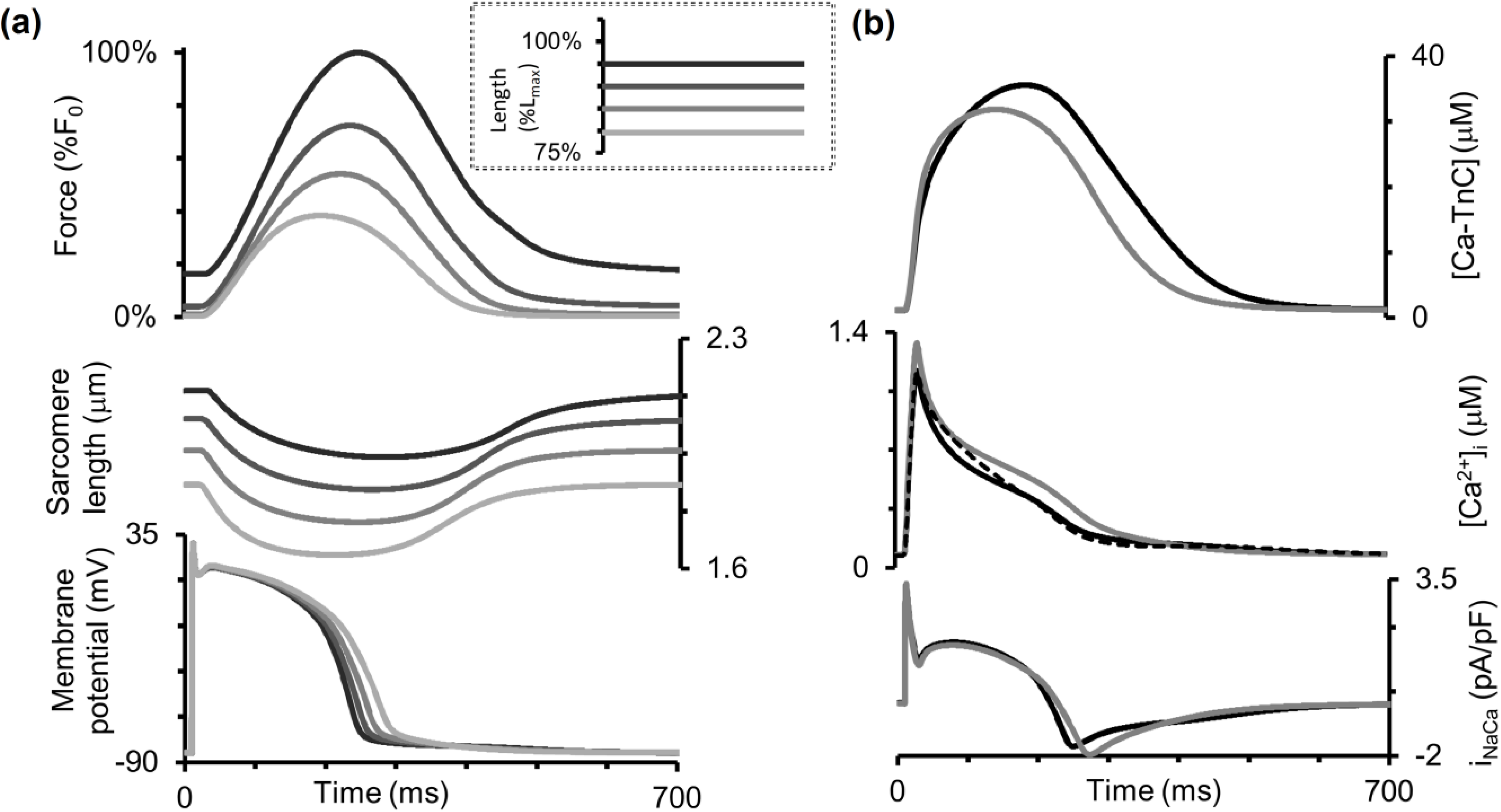
Simulation of isometric contractions at different sample lengths in the TP+M model. The initial length of the sample was decreased from 95% to 80%*L*_*max*_ (see inset from dark to light grey lines). **a** The steady state signals for isometric force (as compared to the force *F*_*0*_ generated at 95%Lmax initial length), sarcomere length, and membrane potential are shown for all lengths considered. **b** The steady state signals for the concentration of calcium-troponin C complexes (*[Ca-TnC])*, intracellular Ca^2+^ concentration (*[Ca^2+^]_i_*) and Na^+^-Ca^2+^ exchange current (*i_NaCa_*) for 85% and 95%*L*_*max*_ initial lengths. Dashed line in the intracellular Ca^2+^ concentration panel is for the numerical experiment where mechano-electric feedbacks were eliminated from the model by imposing isometric conditions on the sarcomere (see text for details)

Note that the AP durations for various lengths in Fig. 4 vary between 230 and 290 ms, i.e. they fall into the experimental range of APDs recorded in a number of experiments on human cardiomyocytes at 37°C with the pacing rate of 1 Hz using microelectrodes [21] (see Fig. 6 of the cited article).

**Fig. 4.**
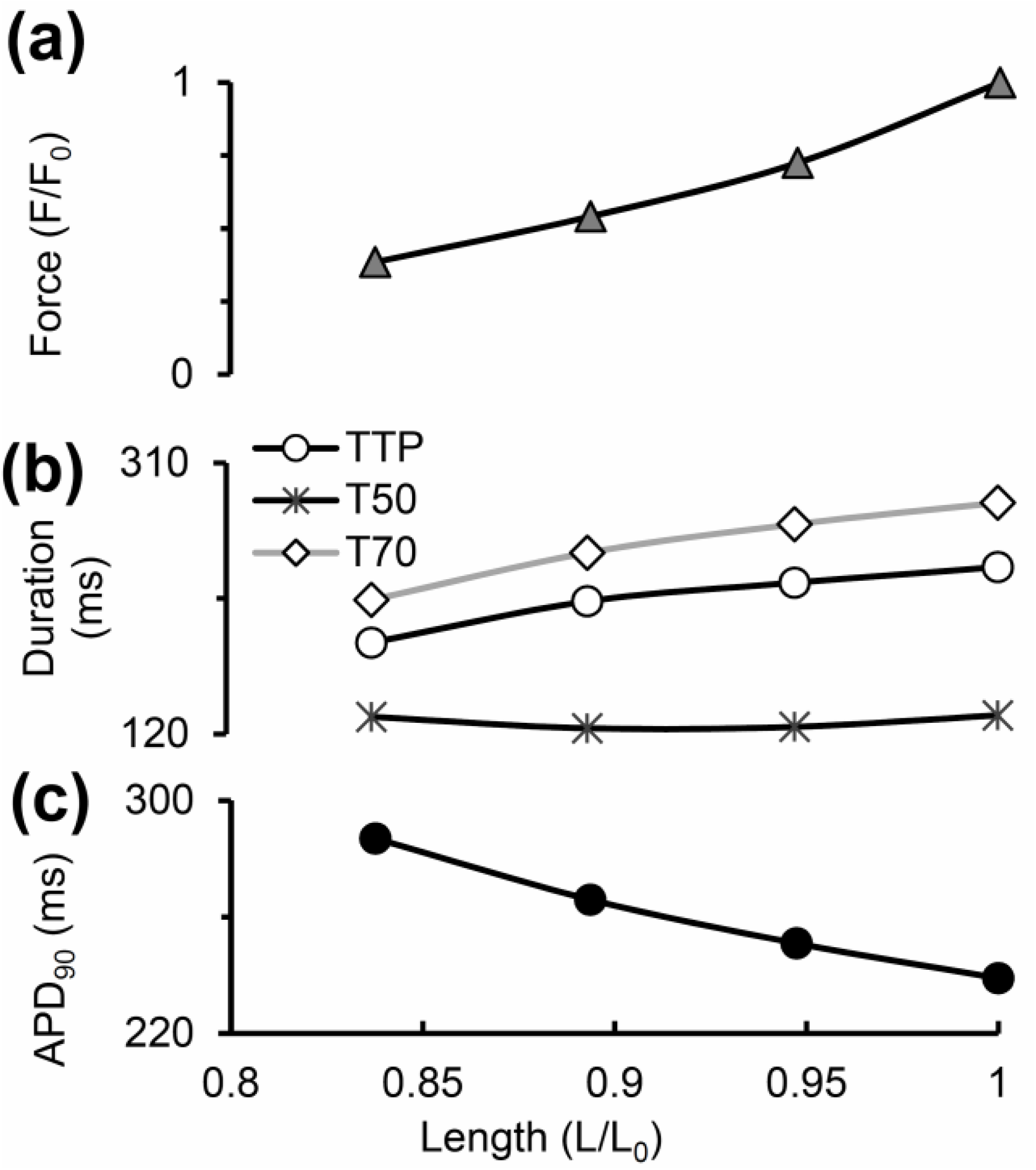
Main characteristics of the isometric cycles in the TP+M model. **a** “Length – Force” diagrams depicting the relationship between length and generated force obtained in a series of isometric contractions with length decreased from reference length *L_0_ = L*_*init*_ (at which the virtual sample generates maximum isometric force *F_0_*) to 84%*L_0_.* **b** Length dependence of isometric twitch temporal characteristics: *TTP* – time to peak twitch; *t*_*50*_, *t*_*70*_ – time to 50% and 70% force decay from peak force in isometric cycles. **c** Length dependence of action potential duration at 90% of repolarization (*APD*_*90*_) in isometric cycles

The temporal characteristics of the Ca^2+^ transient, time to its peak included, are in concordance with experimental data measured in a healthy isolated human ventricular myocyte at 37°C using Fura-2-AM and Fluoforte fluorescence data [21, 60]. It should be noted that the experimental data available on the time-to-peak [Ca^2+^]_i_ in humans are widely varied due, probably, to differences in the dyes and temperatures used and mechanical conditions under which the myocardial preparations were tested. Anyway, it would be impossible to fit the model to all these diverse data simultaneously.

Both time to peak isometric force and isometric relaxation time obtained in our simulations fit well into the range of respective data obtained on papillary muscles, trabeculae and subepicardial biopsy strips from human ventricles [45, 49, 52, 56, 61, 62]. Figure 4b shows the time characteristics of isometric contraction and relaxation for different lengths, namely: time to peak force (*TTP*) and relaxation time, the latter estimated by two indices (*t*_*50*_ and *t*_*70*_). *t*_*50*_ is the time interval from peak contraction (i.e., from *TTP*) to 50% relaxation; while *t*_*70*_, to 70% relaxation. Compared to afterloaded twitches, the mechanics of isometric twitches in the human myocardium has been explored in a large number of experimental studies [42–57]. Although not all of them examined systematically the length effect on the inotropic and lusitropic characteristics of the human myocardium, they have provided us with experimentally observed ranges of TTP, *t*_*70*_ and *t*_*50*_ values for verifying the parameters of the TP+M model that are responsible for its mechanical behavior. Our simulations have been found to be within the ranges of mechanical characteristics obtained in the experiments referred to above.

Figure 4 presents inotropic and lusitropic length dependent characteristics for simulated isometric twitches, in particular for the isometric “Length – Force” relationship (Fig. 4a), which is commonly considered as an isometric equivalent of the Frank-Starling law of the heart for isolated myocardial preparations. The figure plots the peak isometric force against respective length of the virtual sample. Peak force can be seen to increase with length. Such results have been obtained repeatedly for heart muscles from various species and in a number of mathematical models. Here, we repeat this relationship in the TP+M model to make sure that it reproduces correctly this law of the heart.

In contrast to the isometric “Length – Force” curve, the length dependence of calcium and electric signals and characteristics in the simulated series of isometric twitches (see the respective panels in Fig. 3 and 4) should be regarded as model predictions for human cardiomyocytes. Both MCF (*[Ca^2+^]_i_*, Fig. 3b) and MEF (membrane potential, Fig. 3a) are simulated in this series of isometric twitches at various lengths. Fig. 4c reveals a negative dependence of *APD*_*90*_ on length. These simulations together with respective experimental results obtained for other species [63, 64] suggest that the length dependence of both *[Ca*^*2+*^]_*i*_ and APD in isometric twitches may be inherent in real human cardiomyocytes. The mechanisms of this dependence in the model are the same as in the simulation of isotonic afterloaded twitches reported and discussed in detail below.

As in experiments under isometric conditions, in the case of fixed sample length our model accounts for changes in sarcomere length due to the in-series elastic compliance at the preparation boundaries (see Fig. 2). Note that in the model the decay of the Ca^2+^ transient at fixed sample length displays two phases of changing convexity going from positive to negative values and forming a bump. We assumed this biphasic Ca^2+^ decay was related to the change in the sarcomere length during the isometric twitch of the sample. To check this hypothesis, we eliminated the effects of sarcomere shortening in a special numerical experiment by fixing the contractile element, representing sarcomeres in the model (see Fig. 2), at its initial length for the entire twitch. As a result, the Ca^2+^ transient decay in response to isometric sarcomere contraction became monophasic (see the dashed line in the Ca^2+^ transient panel in Fig. 3), thus reproducing the respective feature of the Ca^2+^ transient in the original TP ionic model [20], which does not allow for any mechanical factors. Thus, the biphasic Ca^2+^ decay in the TP+M model is a result of MCF.

In the Discussion below, we will consider experimental studies revealing Ca^2+^ bumps and discuss possible reasons for the variability of this feature in the Ca^2+^ transient in various experiments to the point of its complete absence.

### 3.2 Simulation of afterloaded contractions followed by quick muscle re-stretch to initial length. Model validation

The only experimental study demonstrating MCF in human cardiac muscles (trabeculae) was performed by Vahl et al [55] for very specific afterloaded twitches (Figure 5 here reproduces Fig. 4 from the cited paper). The muscle was allowed to contract against predefined loads, and as the end-systolic length was reached, it was rapidly restretched to its initial length. We performed simulations in the TP+M model using a similar protocol (Fig. 6). For comparison, we can use curves corresponding to an isometric twitch labelled as “0” in Fig. 5 and three afterloaded contractions: one labeled in Fig. 5 as “3” and two adjacent curves (one higher and one lower than “3”).

**Fig. 5.**
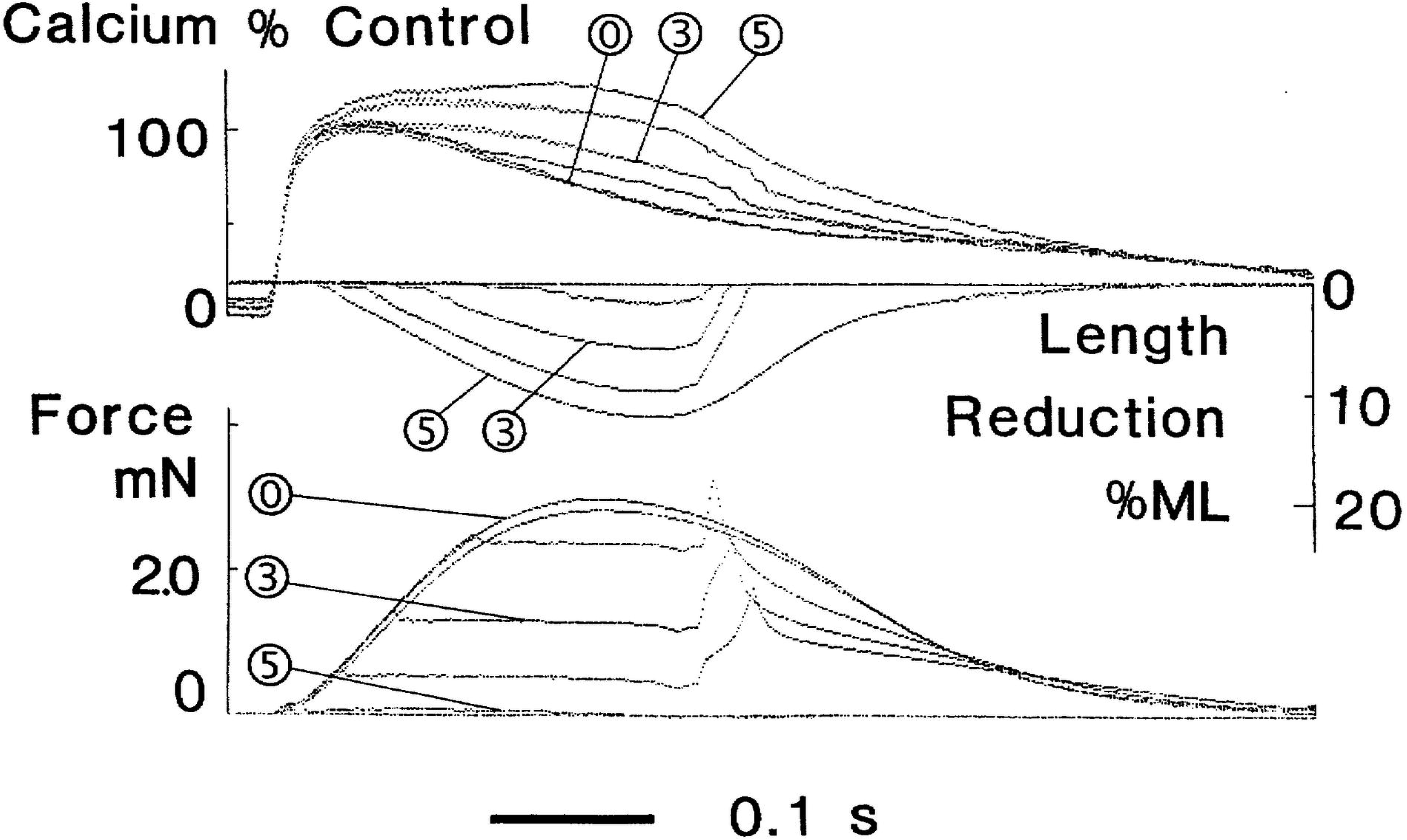
Modified afterloaded contractions recorded in experiments with muscle samples from the left ventricular myocardium of a patient with dilated cardiomyopathy. From top to bottom: intracellular calcium transient, muscle length (ML), and force. The muscle is allowed to contract against predefined loads and as the end-systolic shortening is reached it is restretched at constant velocity to its initial length. “0” indicates isometric contractions; “3”, an afterloaded contraction against an afterload of 40% of maximal isometric force; and “5”, an isotonic contraction against passive resting force. With permission from [55]

**Fig. 6.**
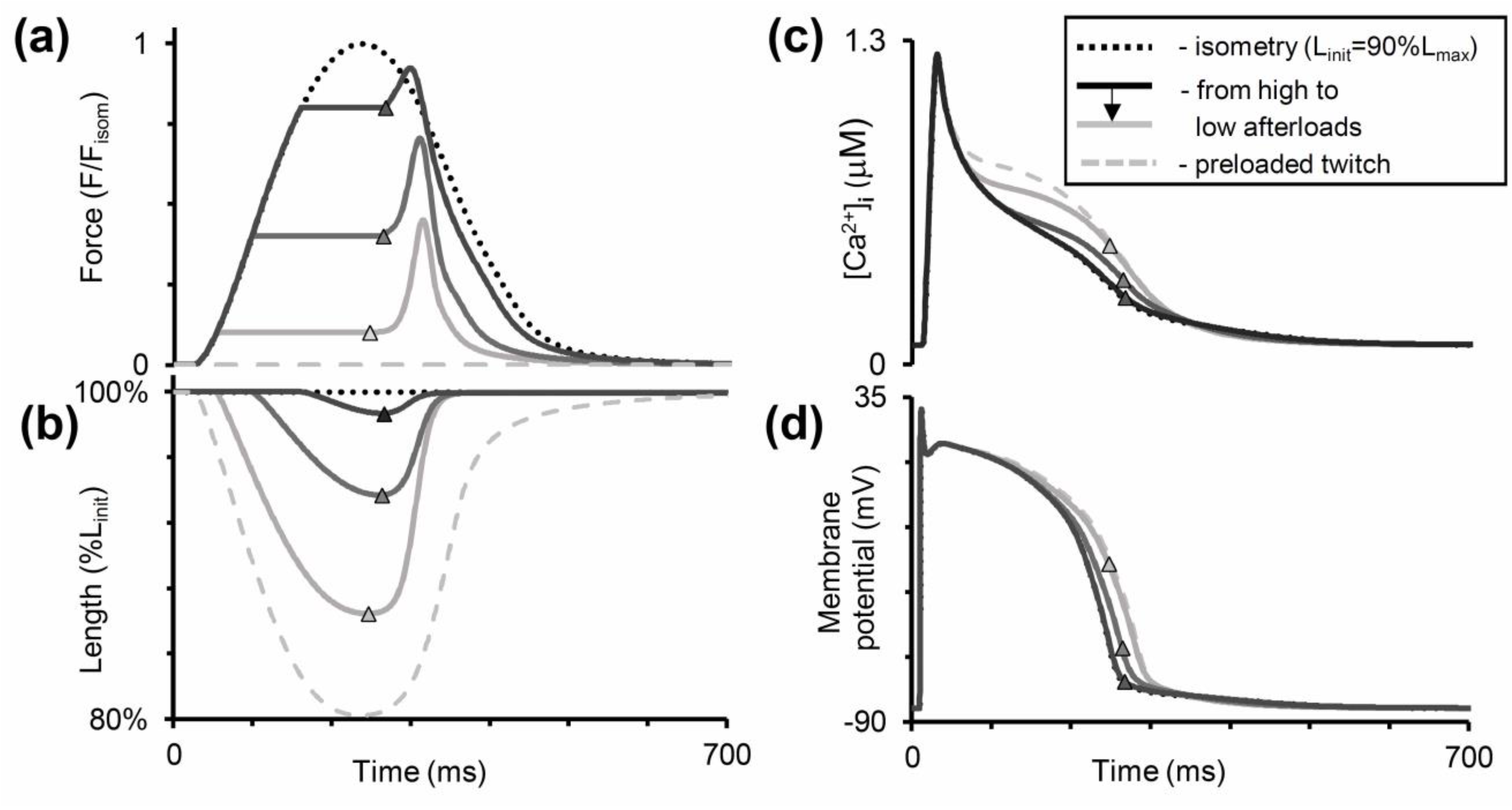
Simulation of modified afterloaded contractions with quick muscle restretching (Fig. 5) in the TP+M model. The mode of cardiac muscle contraction is simulated in the following way. The muscle is allowed to contract against different loads (decreased in **(a)** from dark to light grey lines) in the same manner as in the isotonic afterloaded mode. Then, at the moments of maximum (end-systolic) shortening (shown by triangles) the muscle is forced to stretch at a velocity much higher than that of its lengthening in the full afterloaded cycle (as shown further for the virtual sample in Fig. 7). Thus, the sample quickly returns to its initial length (*L*_*init*_ = 90%*L*_*max*_ in this simulation) and then relaxes isometrically. Dotted lines are for isometric contraction, dashed lines are for preloaded twitch. **a-b** Active force normalized to peak isometric force at *L_init_;* length of the virtual sample (in % of *L_init_*). **c-d** Intracellular Ca^2+^ concentration (*[Ca^2+^]_i_*) and membrane potential

Note that the experimental data in Fig. 5 may be used for model validation with caution since they were obtained on trabeculae from a patient with dilated cardiomyopathy whereas we are trying to verify the TP+M model as that of the healthy human cardiomyocyte. In the same study of Vahl et al [55], Figure 3 shows contractions of a trabecula from a healthy donor heart. Regrettably, for the healthy myocardium these authors presented data on isometric and preloaded twitches only and, therefore, they cannot be used for validating the model in the mode of afterloaded contractions. At the same time, it should be noted that the healthy trabecula in [55] shortened in preloaded twitches twice as much as those from the patient with dilated cardiomyopathy; differences between the Ca^2+^ transients observed under isometric conditions and normal shortening in the healthy trabecula were considerably less than in the cardiomyopathic one. Thus, when comparing the experimental data on afterloaded contractions with the results of our numerical experiment, we deal only with qualitative, rather than quantitative, validation of simulated effects.

Nevertheless, a comparison of the curves in Fig. 5 and Fig. 6 shows that both the real and the virtual preparation become inactivated upon return to the initial length after afterloaded shortening. In other words, in the course of relaxation it cannot develop isometric force to the extent it developed at the same length and over the same time in a full isometric twitch (see the ‘Force’ panels in both figures). Moreover, from the very beginning of muscle shortening and even for some time after restreching to the initial length, afterloaded twitches display a marked surplus of free calcium in the cytosol compared with the isometric twitch in both model and experiment (see the ‘Calcium’ panels in both figures), thus demonstrating MCF. These differences between isometric and afterloaded Ca^2+^ transients cause AP in the model (Fig. 6d) to slow down in the afterloaded twitch. No experimental data are available against which to compare the latter observation and, therefore, this MEF remains a model prediction. The mechanisms that underlie these MCF and MEF in the model are detailed in Section below.

Note that the load-dependence of the Ca^2+^ transients in the model is somewhat different from what is shown in the ‘Calcium’ panel (Fig. 5). Thus, in the model the virtual preparation starts shortening under any afterload following the Ca^2+^ transient peak while in the curves shown in Fig. 5 the beginning of afterloaded shortening precedes it. Therefore the surplus of free cytosolic calcium in afterloaded twitches occurs after this peak in the model and shortly before it in the experiment. This distinction is due to the relatively less steep rise phase of the experimental Ca^2+^ transient compared to the model. We validated Ca^2+^ transients in the model against data from other experiments [21, 60] in which Ca^2+^ transients grew much steeper than in Vahl et al [55]. Therefore the relative positioning of the Ca^2+^ transients in Fig. 6c does not seem impossible. It is noteworthy that Fig. 5 shows data characteristic of dilated cardiomyopathy. Aren’t the relatively slow rise phase of the Ca^2+^ transient and related earlier occurring (before the Ca^2+^ transient peak) surplus of free cytosolic calcium manifestations of a specific pathology? This supposition appears to be reasonable if we consider Fig. 4 from another paper of the same authors [65]. According to this figure, in the case of mitral valve stenosis the Ca^2+^ transients peaks were similar under isometric conditions and in a preloaded twitch while after the peak the preloaded twitch produced a noticeable surplus of free cytosolic calcium compared to the isometric conditions.

In the ‘Discussion’, we will consider and additionally discuss one more experimental result revealing a higher peak Ca^2+^ transient recorded for a preloaded twitch as compared to an isometric one in rat myocardium instead of human one [66].

### 3.3 Simulation of isotonic afterloaded twitches

The effect of load dependent relaxation in isotonic afterloaded twitches most clearly demonstrates the influence of mechanical conditions of myocardium contraction and relaxation on its activation/inactivation. This effect was demonstrated for the first time more than forty years ago in experiments on mammalian and frog papillary muscles and trabeculae, which revealed higher muscle lengthening (relaxation) velocities in isotonic afterloaded twitches under smaller loads than larger ones [67, 68]. This paradoxical effect pointed to the inactivation of muscle contractility in response to the reduction of the applied load. Later this effect was confirmed in experimental studies on various mammals – rat, rabbit, cat, etc. [41, 58, 59].

Figure 7 shows the results of our simulations using the TP+M model for a steady-state isometric contraction of the sample initially stretched by a preload to a length of 90%*L*_*max*_ (where *L*_*max*_ corresponds to the sarcomere length equal to 2.23 μm). Both time to peak isometric force and isometric relaxation time obtained in our simulations fit well into the range of respective data obtained on papillary muscles, trabeculae and subepicardial biopsy strips from human ventricles [45, 49, 52, 56, 61, 62]. The isometric signals are shown together with ones recorded in simulated isotonic afterloaded contractions under various afterloads, at the same initial length of 90% *L*_*max*_.

**Fig. 7.**
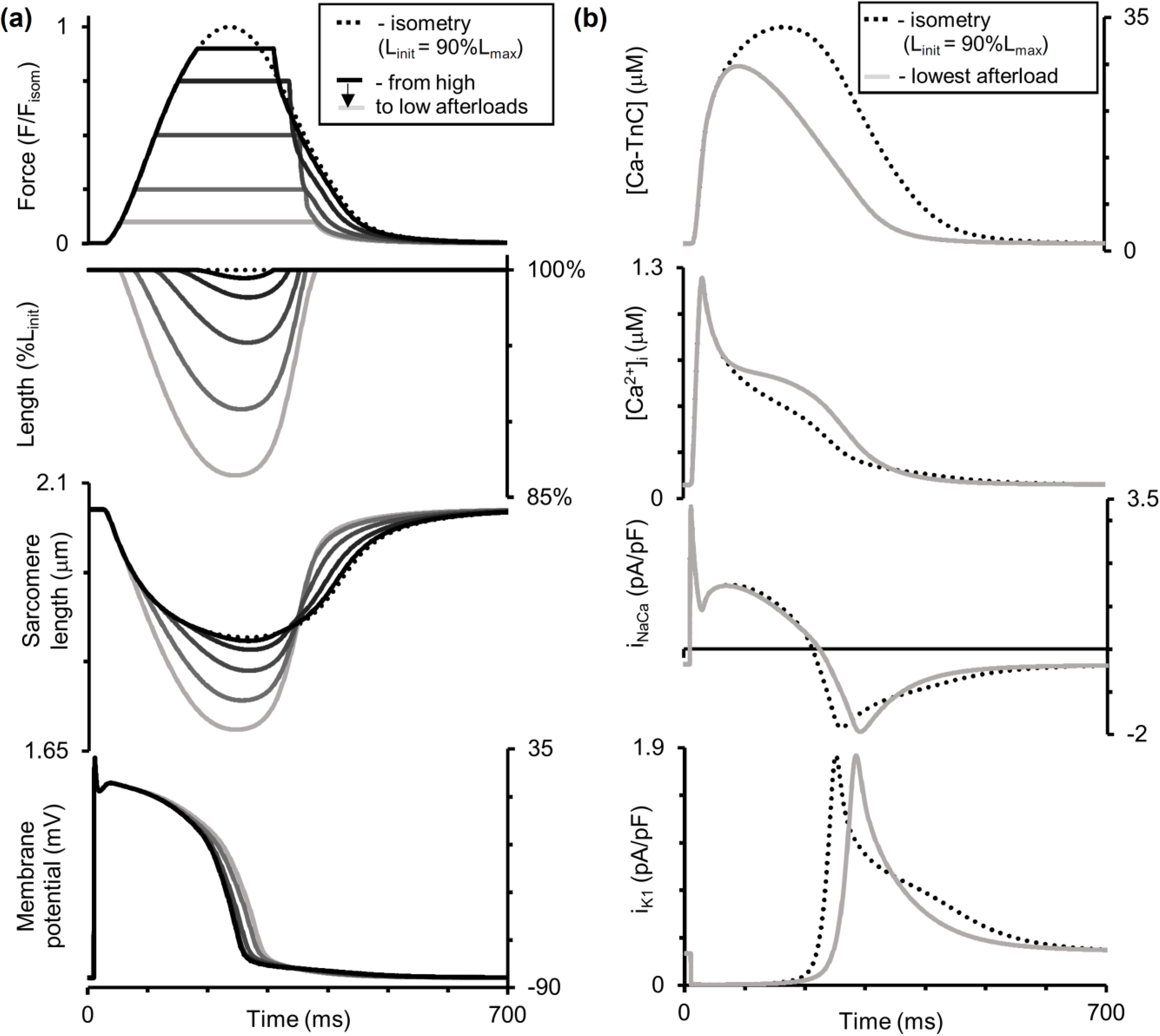
Simulation of a series of isotonic afterloaded contractions in the TP+M model. The initial virtual sample length *L*_*init*_ is equal to 90%*L*_*max*_. *F*_*isom*_ is a peak active isometric force at *L_init_*. Each afterload was applied following a steady-state isometric contraction (dotted lines) varying from a high afterload of 0.9*F*_*isom*_ (black lines) to a low afterload of 0.1*F*_*isom*_ (light grey lines). **a** Time-dependent signals of the generated force (*F/F_isom_*), sample length, sarcomere length and membrane potential for various loads applied. **b** Time-dependent signals of the concentration of calcium-troponin C complexes (*[Ca-TnC]*), intracellular Ca^2+^ transient (*[Ca^2+^]_i_*), Na^+^-Ca^2+^ exchange current (*i_NaCa_*) and inward rectifier K^+^ current (*i_K1_*) for isometric and afterloaded contractions at low afterload

Figure 7 demonstrates load dependent relaxation in the TP+M model, i.e. an increase in the velocity of muscle lengthening (isotonic relaxation) with a decrease in the applied load (see the length panel in Fig. 7). As a result, the smaller the afterload, the greater the difference between the duration of the isotonic of shortening-lengthening phase and that of the phase of an isometric twitch during which the generated force is higher than the applied afterload. The isometric and isotonic afterloaded forces shown in Fig. 7a reveal these differences. To quantify this difference, the load dependence index (LDI) is calculated at any certain level of afterload *F*_*isot*_ as the ratio of the time intervals shown in Fig. 8: LDI = *t*_*isot*_/*t*_*isom*_ [59].

**Fig. 8.**
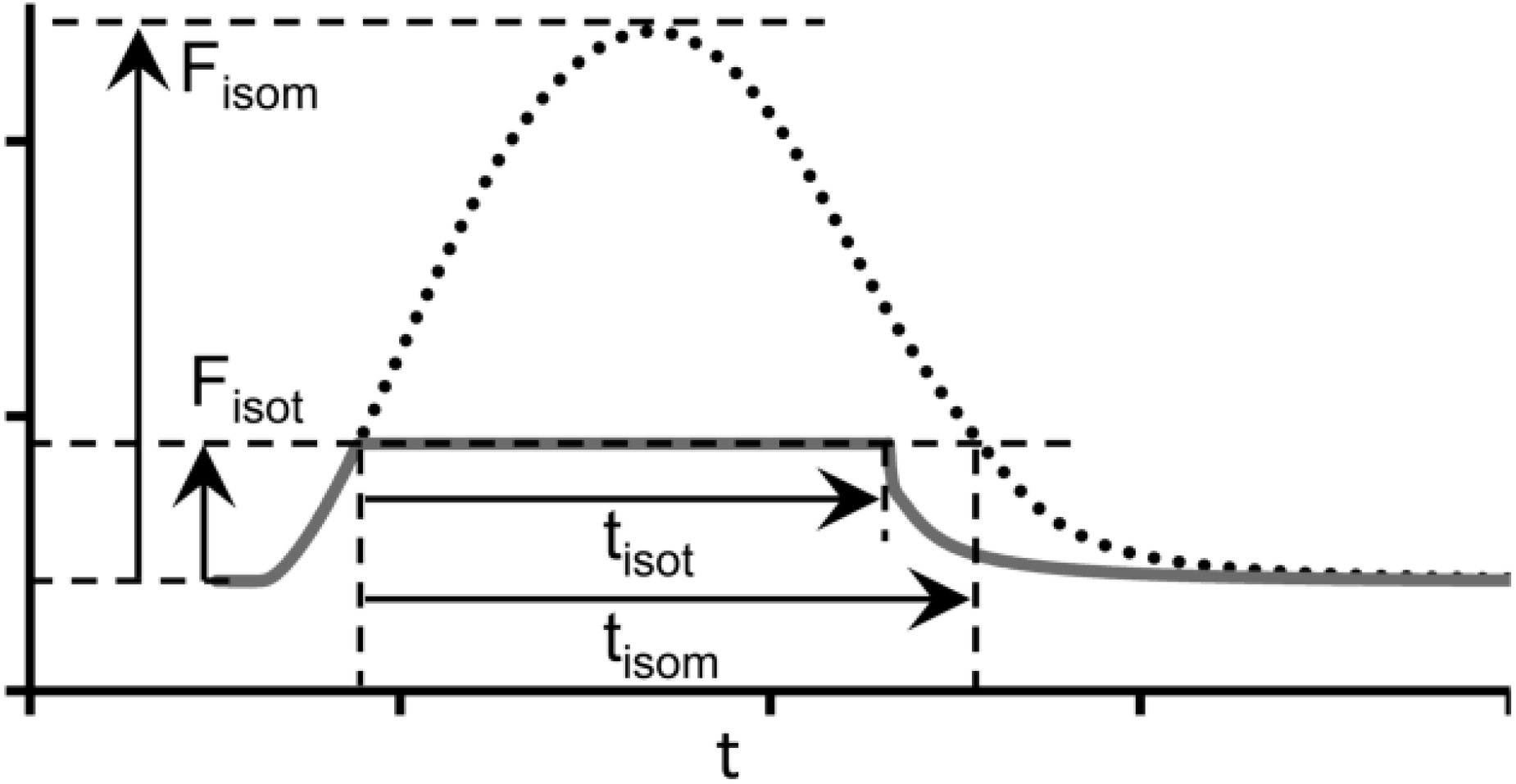
The load dependence index (*LDI*) is set as the ratio *t*_*isot*_*/t*_*isom*_

Figure 9a shows the dependence of LDI on the normalized afterload *F_isot_*/*F*_*isom*_ at two initial lengths (90%*L*_*max*_ and 95%*L*_*max*_) in the TP+M model. For both initial lengths, LDI is seen to lie lower than 1 at medium and low afterloads reflecting the effects of muscle inactivation by shortening in the model. This model prediction for the human myocardium fits well with the data obtained on various species, e.g. guinea pigs [59] and rats [69]. Unexpectedly, the LDI curve is independent of the initial sample length (preload) of the virtual sample.

**Fig. 9.**
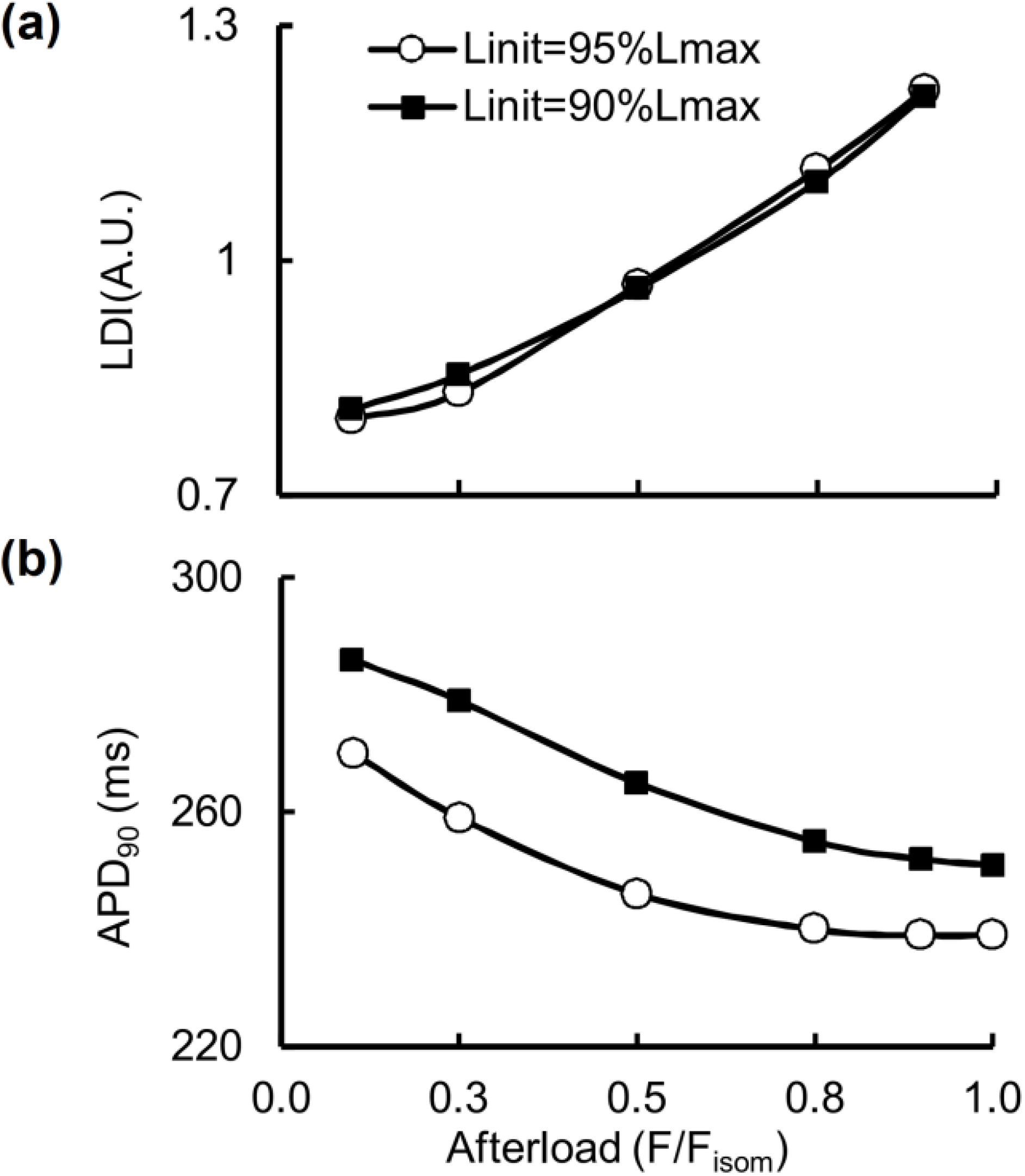
Load dependent characteristics in the TP+M model for two initial virtual sample lengths *L*_*init*_: 90% and 95%*L*_*max*_. The abscissa plots the force values normalized to the maximum isometric value *F*_*isom*_ corresponding to the initial length *L*_*init*_. **a** The load dependence index *(LDI)* calculated as shown in Fig. 8. **b** Action potential duration calculated at 90% repolarization (*APD*_*90*_)

Figure 7b shows calcium transients (*[Ca^2+^]_i_*) in the virtual muscle cell during isometric (dashed) and isotonic afterloaded (solid) contractions at low load. The intermediate traces for Ca^2+^ transients lie between these curves, but they are not shown for simplicity. The load-dependent slowing of the Ca^2+^ transient decay phase was shown to be present in mammalian myocardial preparations [58]. This MCF effect arises in the TP+M model due to increased dissociation of Ca-TnC complexes (*[Ca-TnC]*, Fig. 3b) in response to faster and greater shortening under low load (see the changes in sample and sarcomere length in Fig. 7b). The decreased dissociation occurs as a result of the length dependent cooperativity mechanisms in myofilaments.

This MCF, in turn, induces MEF manifestations in the model as shown in Fig. 7a: the smaller the load, the slower the repolarization phase and the greater the AP duration (APD). Similar load dependent increase in APD was shown in cardiac preparations from mammals [58].

Note that the combined model with the baseline TP parameters of Ca^2+^ currents did not produce such an AP response to the shortening, showing almost no effect of the mechanical load on AP. In order to simulate the load dependence of AP in the TP+M model, we performed additional tuning of the parameters for the NCX current (iNaCa, Fig. 7b). This current had been shown to trigger AP modulations in response to load dependent modulations of the cytosolic Ca^2+^ concentration in the ‘Ekaterinburg – Oxford’ model [9]. An analysis of the TP+M model showed that NCX is able to “convert” a higher cytosolic Ca^2+^ concentration into slowing repolarization at low load only if the reverse mode NCX current contributes essentially to the repolarization process. During the reverse mode, one Ca^2+^ ion comes from the extracellular space into the cytosol in exchange for three Na^+^ ions coming out from the cell. The higher the cytosolic Ca^2+^ concentration (i.e. the smaller the transmembrane Ca^2+^ gradient), the smaller the NCX current during the reverse mode. In the original TP model, this NCX phase during the AP plateau was absent. To make the model produce the reverse mode NCX current, we changed the value of only one parameter in the NCX equation (4). This parameter α (see the Supplement for NCX current description) controls the ratio between the positive and negative terms in the NCX formulation and thus allows one to obtain a positive outward NCX current during the AP plateau, and this outward NCX current contributes to AP repolarization.

It should be pointed out that our assumption of the occurrence of a reverse NCX current during the cardiac cycle is speculative, but similarly speculative is the opposite assumption of its absence made in the original TP model. Because there are no direct experimental data on the NCX time course in human cardiomyocytes, both assumptions can only be justified by speculations on the physiological feasibility of the reverse mode for cellular outputs. In terms of AP generation, the outward NCX current during the AP plateau phase counteracts the inward L-type Ca^2+^ current and thus does not help maintain the plateau (e.g. see paper [70]). In ionic models focused only on AP description, this reason can possibly be employed to exclude the reverse mode NCX current without the loss of accuracy in simulated AP. In terms of contraction, however, which is the main function of the cardiomyocytes, the reverse NCX current brings Ca^2+^ into the cell, thus co-acting in the same direction with the L-type Ca^2+^ current and allowing more Ca^2+^ in to support myofilament activation and further contraction. Importantly, the latter considerations were formulated in an experimental work dealing with the reverse NCX current in human ventricular myocytes [71].

Given the arguments put forward by Weisser-Thomas et al. [71] and the important role of the reverse NCX current for MEF in the cardiomyocytes of most mammals, we assume that the reverse mode NCX does exist.

As we previously showed and analyzed it in detail by the ‘Ekaterinburg – Oxford’ model, NCX is the primary mechanical sensor for Ca^2+^ dissociation from Ca-TnC complexes [9, 72]. It works only as a trigger of AP changes. The secondary effect is a change in the time course of the voltage-dependent currents induced by the initial shift triggered in AP.

For analyzing primary and secondary mechanically induced events we used the ‘Difference-Current Integral Method’ described in detail earlier [72]. This method allows one to take into account all the relative instant deviations of each current in the afterload twitch from the same current's trace during the isometry and to evaluate the total contribution of these deviations to the difference between the afterloaded and isometric APs. Analysis by this method revealed that for smaller afterloads NCX current is the first which changes begin to deviate AP trace immediately upon switching from the isometric loading mode to the afterloaded one (54 ms; Fig. 7b, *i*_*NaCa*_). As a result, for smaller afterloads the action potential achieves a higher level at the beginning of the plateau (Fig. 7a), thus enhancing voltage-dependent relative changes in the inward rectifier K^+^ current (Fig. 7b, *i*_K1_). Relative changes in *i*_*K1*_ tended to prolong AP are so pronounced that oppositely directed relative changes in some calcium or/and voltage-dependent currents (*i*_*CaL*_, *i*_*Ks*_, *i*_*pK*_; not shown) can no longer seriously influence the slowing of AP prolongation. Even the inversion of the current difference between the afterloaded NCX current and the isometric one (reversal of the sign of this difference from “−” to “+”) after ~210 ms, which therefore begins to contribute to the AP shortening after this moment, cannot overcome the relative changes in the inward rectifier K^+^ current. The change in the voltage-dependent current *i*_*Kr*_ relatively to isometric time course works to AP prolongation as well as *i*_*K1*_ current, but it is less pronounced than the latter.

Figure 9b presents the dependence of *APD*_*90*_ on the relative afterload for isotonic afterloaded twitches at two initial lengths of the virtual sample: 90%*L*_*max*_ and 95%*L*_*max*_. For each initial length, *APD*_*90*_ can be seen to increase with a decrease in the afterload. The AP becomes longer by up to 14% and up to 13% compared to the isometric conditions for 90%*L*_*init*_ and 95%*L*_*init*_, respectively (Fig. 9b). It can also be seen that *APD*_*90*_ depends significantly on the initial length: *APD*_90_ for 90%*L*_*max*_ is located much higher than for 95%*L*_*max*_. By contrast, “Afterload – LDI” does not depend on the initial length (Fig. 9a). This significant distinction between the “Afterload – APD_90_” curves for different preloads (initial lengths) is a model prediction and, as far as we know, this effect has not yet been studied in experiment either for human or for animal myocardium.

In the 1950s, the slope of the isometric “Length – Force” curve was attributed solely to changes in the length of the zone of overlap between thick and thin filaments. As more careful measurements were made, it became clear that this slope is steeper than one produced only by changing the length of the overlap zone. Later, calcium inactivation of the contraction during sarcomere shortening (i.e. MCF) was generally recognized as also contributing to the slope of the isometric curve. For comparison, Fig. 10 shows an isotonic (afterloaded) “Length – Force” curve (solid line) for the initial length 90%*L*_*max*_, together with a respective fragment of the isometric “Length – Force” curve (dashed line) obtained in the TP+M model. The steeper slope of the isotonic (afterloaded) curve is known from numerous experiments on animals [41]. The TP+M model predicts the same for the human myocardium. The steeper slope of the isotonic curve is due to the higher sarcomere shortening velocity in isotony compared to isometry, since the kinetics of cross-bridge attachment/detachment in the model is regulated by not only the length but also velocity of sarcomere shortening/lengthening. Increasing the velocity decreases the number of attached cross-bridges (via the cooperativity mechanisms) and thus facilitates the inactivation of Ca-TnC complexes.

**Fig. 10.**
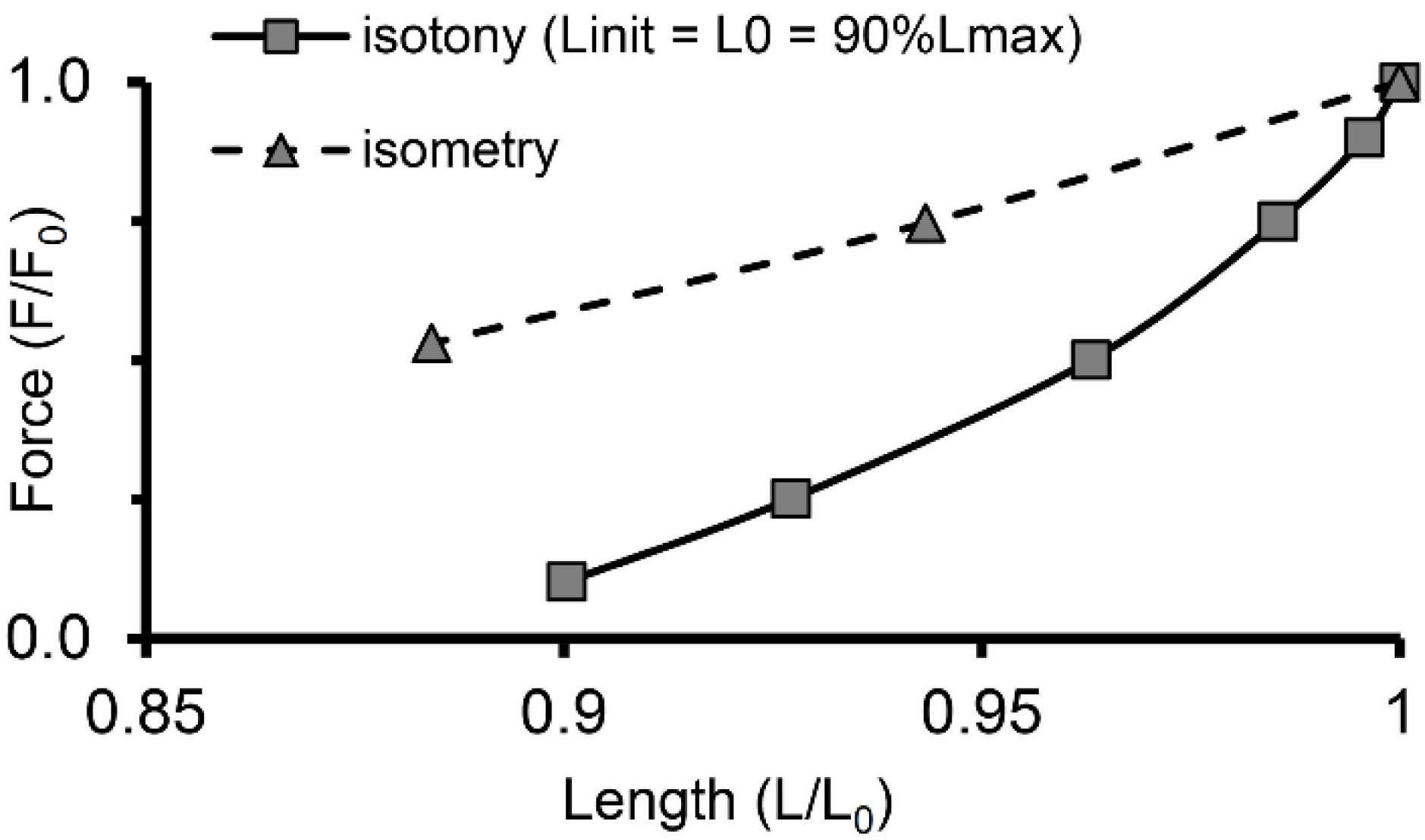
“Length – Force” diagram depicting relationship between end-systolic shortening and end-systolic force obtained in a series of isotonic afterloaded contractions at initial length *L*_*init*_ = 90%*L*_*max*_ (solid line) compared with relationship between length and generated force obtained in a series of isometric contractions (dashed line), where length decreases from reference length *L*_*0*_ = *L*_*init*_ (at which the virtual sample generates maximum isometric force *F*_*0*_) to 88%*L*_*0*_

### 3.4 Simulation of modified isotonic afterloaded twitches with quick load switches

Figure 11 shows the results of numerical experiments in which the load on the virtual sample was changed abruptly during isotonic afterloaded contraction-relaxation cycles. The initial length was 90%*L*_*max*_ and the load was either increased (Fig. 11a) or decreased (Fig. 11b). In both cases, load switching occurred approximately in the middle of the initial isotonic cycle, i.e. when the isotonic shortening phase under the first load was nearly at its end. Such experiments had been performed previously involving heart muscles from guinea pigs [59] and cats and rabbits [41], and mechanical responses to load switching qualitatively similar to the ones shown here (*Force, Length*, Fig. 11a-b).

**Fig. 11.**
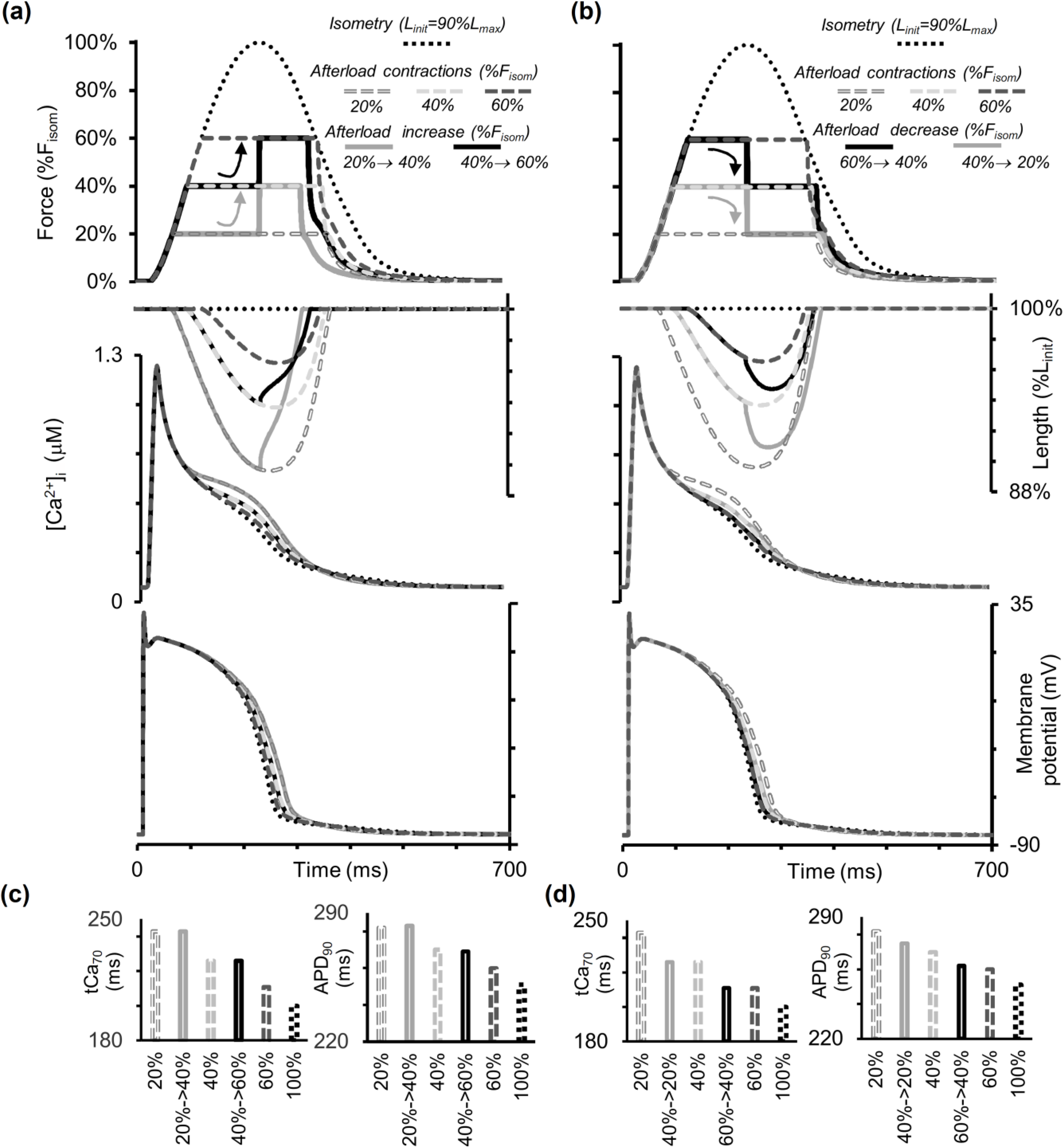
Simulation of a quick change in the load during an isotonic afterloaded cycle in the TP+M model. The initial virtual sample length *L*_*init*_ is equal to 90%*L*_*max*_. Dotted lines are for isometric contraction. Dashed lines show afterloaded contractions under different loads (20, 40 and 60 % of peak isometric force *F*_*isom*_ at *L_init_*). Solid lines show the traces for quick increase **(a)** and decrease **(b)** in the load approximately in the middle of the isotonic plateau. From top to bottom in both **(a)** and **(b)** : force, sample length, intracellular Ca^2+^ transient (*[Ca^2+^]_i_*) and membrane potential changes. **c-d** The value *tCa*_*70*_ is time from peak Ca^2+^ transient to 70% decay. *APD*_*90*_ is action potential duration at 90% repolarization

The corresponding responses of the calcium transients (*[Ca^2+^]_i_*) and membrane potentials to the changing of the applied load are shown in Fig. 11a-b. The diagrams in Fig. 11c-d show the *APD*_*90*_ values and the duration of the calcium transients (*tCa*_*70*_) for all switchings applied. Despite major changes in the mechanical conditions during the cycle, we do not observe any significant changes in the characteristics of the APs and Ca^2+^ transients compared with their values in the simulated usual afterloaded isotonic twitches under the first load. This simulation result suggests that mechanical interventions imposed during the contraction phase affect Ca^2+^ kinetics and electrical activity more significantly than interventions imposed during the relaxation phase.

### 3.5 Simulation of mechano-calcium and mechano-electric feedbacks in response to quick release of the human myocardium at different moments during the isometric twitch

Figure 12 shows the results of instantaneously releasing the virtual sample, initially contracting isometrically at the length of 90%*L*_*max*_. The sample shortens quickly by 5% of the initial length at different moments during the isometric twitch (Fig. 12a) and then contracts/relaxes in the isometric mode at this shorter length (Fig. 12b). Similar experiments had been performed on papillary muscles from various animals, for example, cats and ferrets [58, 73].

**Fig. 12.**
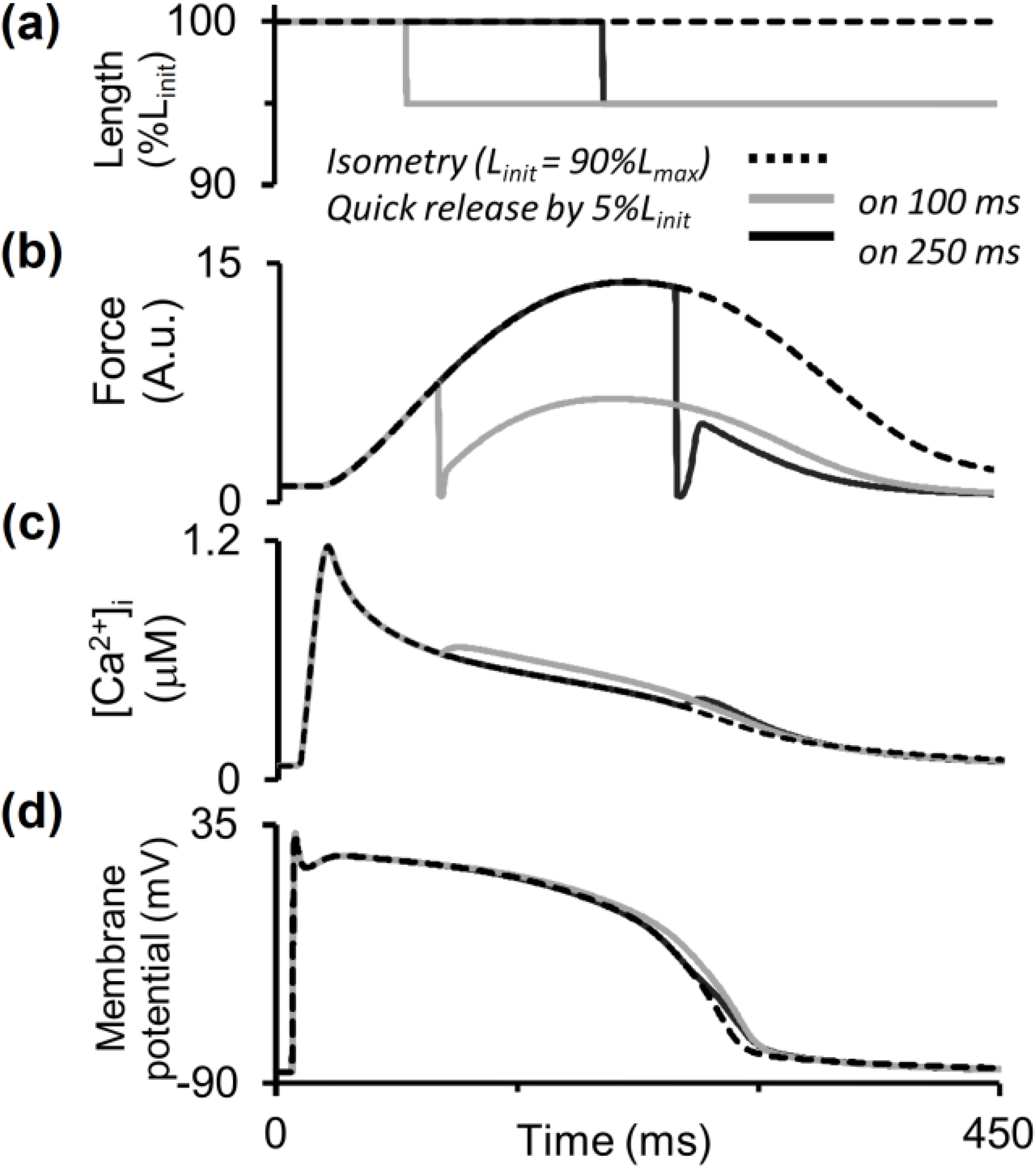
Simulation of quick releases of the virtual sample during the isometric cycle in the TP+M model. The initial length *L*_*init*_ is equal to 90%*L*_*max*_. The sample is quickly released to 95%*L*_*init*_ at 90 and 240 ms after the stimulus and is held at a new length until the end of the cycle. **a-d** Virtual sample length, force development, intracellular Ca^2+^ transient (*[Ca^2+^]_i_*) and membrane potential in the simulation

Mechanical inactivation manifests itself as a decrease in force developed by the virtual sample accompanied by an increase in free cytosolic calcium during the decay phase of the Ca^2+^ transient (Fig. 12c) and by AP prolongation (Fig. 12d). Additional free cytosolic Ca^2+^ arises after the quick release due to additional Ca-TnC dissociation caused by the mechanical perturbation. As well as in the simulations presented in Fig. 7, here we also observe APD prolongation (Fig. 12). However, the mechanism of this effect is different. In Fig. 7, the prolongation was a result of reduced repolarization current produced by NCX in the reverse mode. In the case of the later length release in Fig. 12, the reverse NCX current has already turned into forward depolarizing current. In the model, therefore, the mechano-dependent increase in free cytosolic Ca^2+^ concentration enhanced this depolarizing NCX current and caused the AP prolongation observed. Thus, an abrupt decrease in length resulted in the prolongation of both calcium transient and AP even during the relaxation phase.

The TP+M model also provides evidence for the dependence of Ca^2+^ transient and AP time course on quick release amplitude (not shown).

## 4 Discussion

MEF and MCF are an important way for cardiomyocytes to adapt to mechanical conditions under which they contract in the heart chambers walls, including conditions of mechanical interaction between heterogeneous cardiomyocytes. Elucidating the mechanisms that underlie MEF and MCF presents one of the most complicated physiological problems. A number of studies, including experimental works and modeling, have been carried out to shed light on this subject [1–3, 9, 24, 74–76]. MEF and MCF effects can be divided into two large groups: fast and slow responses of the cardiomyocyte to mechanical impacts. Each of these two groups requires a separate study, since there are serious grounds to believe that the mechanisms underlying them may not be the same.

In particular, it is generally recognized that the contribution of such direct MEF sources as stretch-activated channels (SACs) is a very important factor for slow responses. However, explaining the fast responses that take place during one contraction-relaxation cycle requires involving other, more complex multi-step mechanisms.

To study the fast responses of the human cardiomyocyte to mechanical impacts, we developed a TP+M model of electromechanical coupling in the human cardiomyocyte based on the ‘ten Tusscher – Panfilov’ electrophysiological model and the mechanical activity module taken from the ‘Ekaterinburg – Oxford’ model. We implemented the main experimental modes of the cardiac muscle contraction-relaxation cycles (isometric and isotonic afterloaded ones) in this TP+M model and studied the effects of mechanical conditions on the contractile cycle.

The TP+M model was used to investigate the fast responses of the virtual heart muscle to the mechanical modes as follows:

– responses to loads under which the sample contracts in the isotonic afterloaded mode and in various modifications of the latter (Fig. 7, 11, 6);
– responses to the fast shortening of the sample at different moments during the contraction-relaxation cycle (Fig. 12).

Releasing the heart muscle quickly to a shorter length (Fig. 12) within just a few milliseconds reduces the mechanical activity of cardiomyocytes throughout the contraction-relaxation cycle much more significantly than what could be achieved by simply decreasing the thin and thick filament overlap zone in the sarcomeres. Below we will discuss the reasons why SACs would be irrelevant for explaining this effect.

The only part of our present study where we deviated from the subject of fast responses was concerned with the simulation of isometric contractions from various initial lengths (pre-stretching) of the virtual sample. The reason for the deviation was that we used in the study the results of these simulations to verify the model by comparing these results with the corresponding data from physiological experiments [56]. These physiological data had been obtained in a series of steady state isometric contractions with a fixed pacing rate of 1 Hz at different lengths of each tested muscle preparation. Accordingly, the model was put in the same conditions. Thus, the characteristics of isometric contractions shown in Fig. 3 and Fig. 4 were obtained at different lengths just in the simulation of the series of steady state isometric contractions at 1 Hz. These characteristics indeed fall into the ranges of values reported in both the cited work [56] and other experimental studies making available data for isometric contractions of the human myocardium at one or another length, although the they did not specially investigated the effect of differences in length [45, 51, 52, 62].

According to the TP+M model, mechanical conditions should have an effect on the duration of both action potential and calcium transients in the human cardiomyocyte and contribute substantially to their mechanical activity. According to the simulation, the smaller the afterload in isotonic cycles, the longer the duration of AP and Ca^2+^ transient and the shorter the duration of the mechanical twitch (Fig. 7). These results are TP+M model predictions, which currently cannot be directly experimentally confirmed or rejected due to the lack of respective experimental data for the human myocardium. Nonetheless, these modeling results seem to be plausible since similar MEF and MCF manifestations have been found in numerous experiments on animals.

Indeed, a large number of classical experimental studies involving heart muscle preparations [58, 63, 73, 77] and theoretical explorations with the help of mathematical models [9, 24, 78, 79] have been carried out to identify the mechanisms that determine fast myocardium adjustment to changes in contraction mechanics, particularly in length or applied load. Max Lab has demonstrated in his experiments that additional free calcium arises in the cytosol in response to a decrease in length or load. This surplus of the calcium transient is partially retained during most of the subsequent phase of the contraction-relaxation cycle, even if the mechanical impact has already ceased to act (for example, if, after a quick release, the preparation has been quickly returned to its initial length) [58]. In the same experiments, the AP duration was found to increase along with this modulation of the calcium transient, while the ability of the muscle to develop active force decreased, and this mechanical inactivation continued until the very end of the contraction-relaxation cycle.

As was mentioned above, these load and length dependent events cannot be explained by SACs because, firstly, additional free calcium arose in response to shortening rather than stretching; secondly, if this additional Ca^2+^ initiated by the mechanical impact entered the cell from the outside, then it would sooner activate the cell additionally whereas in reality the cell was deactivated. Many of the above-cited studies suggested the following mechanism to explain all of these events together. As the sarcomere shortens, the dissociation of Ca-TnC complexes increases causing both observed events to happen simultaneously: inactivation and increasing free Ca^2+^ concentration. This additional free cytosolic Ca^2+^, in turn, increases AP duration via the following chain of events. The surplus of free cytosolic Ca^2+^ causes a slight decrease in the NCX current. At this moment, NCX acts in the reverse mode (for example, see *i*_*NaCa*_ in Fig. 7b). Thus, its decrease results in a slight decrease in the total repolarizing (outward) currents. Despite being apparently small, this effect starts to slow down the AP repolarization. It initiates a delay in the activation of the potassium currents (see Fig. 7b for *i_K1_*), followed by APD prolongation.

In this hypothetical chain of intracellular mechanisms, the most difficult to explain was the assumption of increased Ca-TnC dissociation in response to myocyte shortening. We first proposed a possible explanation for this increase and justified it by mathematical modeling in 1991 [41]. The key point of that proposal was one of the known mechanisms of cooperative interaction between regulatory and contractile proteins: the lower the concentration of cross-bridges attached to the thin filaments near a Ca-TnC complex, the more probable the dissociation of this complex [13, 80, 81]. Combined with the decreasing probability of cross-bridge attachment during sarcomere shortening, this cooperativity provided a clue to understanding the length dependent decrease in the Ca-TnC decay. What particular mechanisms underlie the effect of length on cross-bridge attachment is currently a matter of much debate [35, 38, 82, 83]. For a long time, the most popular explanation for this effect was the concept of lattice spacing that related the sarcomere interfilamentous distance and, thus, the probability of cross-bridge attachment to sarcomere instant length [35]. However, recent experimental studies have suggested that it is the giant protein titin rather than the lattice spacing effects that determines the length dependence of cross-bridge attachment [38]. It has also been conjectured that lattice spacing modulation does exist, but it is titin-based [83]. Anyway, length dependence is invariant with regard to its underlying mechanisms and is accepted by all authors.

The incorporation of these mechanisms first into the model of myocardial mechanical activity [41] and then into the ‘Ekaterinburg – Oxford’ model of electromechanical coupling in guinea pig and rabbit cardiac muscles [9] allowed us to reproduce and explain practically the whole range of phenomena associated with fast responses to mechanical impacts within the framework of these models.

Later, other authors used similar approaches in their models, for example, Landsberg and Sideman [84], Jeremy Rice [26], and Lauren Dupuis [85, 86]. These authors studied the contribution of mechanical conditions of contraction to myocardial mechanical activity and applied their models to assess and analyze *in vitro* data obtained in experiments on various warm-blooded animal heart muscles. The contribution of length and/or load to the Ca-TnC kinetics is justified differently in these models and is, consequently, represented by different formulas in the equations describing this kinetics.

Thus, for example, the Dupuis model, like ours, considers the cooperative effect of cross-bridges on the affinity of Ca-TnC complexes to be the driver of the Ca-TnC mechano-dependent kinetics. Moreover, the authors of this model substantiate very carefully the formulas that determine the contribution of this type of cooperativity. As a result, the model reproduces adequately both the load dependence of isotonic afterloaded twitches and the length dependence of isometric ones. The calcium transient in the original Dupuis model is defined simplistically as an explicit function of time; therefore, the problem of MCF and MEF cannot be investigated directly within the framework of this model. To be able to solve it, the Dupuis model should be combined with one or another model of intracellular Ca^2+^ handling and AP generation, and the result will obviously depend on the properties of all components combined.

In the Rice model, the contribution of length to the Ca-TnC kinetics is represented by a rather arbitrary simplified formula, and this model reproduces successfully the length dependence of isometric twitches (including the influence of preloads, i.e. lengths, on the Ca^2+^ transients in these twitches), but not the load dependence of isotonic twitches.

Despite the repeatedly confirmed presence of fast MEF and MCF effects in the myocardium of various animals (rat, cat, rabbit, dog, guinea pig, etc.), it still remains unclear whether these effects are characteristic of the human myocardium, and even if they are, whether the mechanisms that underlie them are the same. The availability of only limited experimental data has been the reason why there are still no clear answers to these questions within the framework of experimental physiological studies. In this context, it is becoming essential to use mathematical modeling as a tool for at least tentative prediction of possible answers. It is noteworthy that the possible answers to the above-stated questions proposed within the framework of the model-based approach were *a priori* far from being obvious. For instance, in a recent attempt to assess MEF effects in the framework of an electromechanical model formed by coupling the Rice mechanical model with electric models for various animal cardiomyocytes, it was concluded that MEF effects occur in the canine myocardium but they are practically absent in the human cardiac muscle [24]. It is clear that the results obtained by these authors depended significantly on the properties of the models employed, including those of the Rice model we just mentioned, and, therefore, their conclusion seems quite debatable, especially considering the presence of such feedbacks in all other studied mammals.

Admittedly, although the above-described MCF and MEF mechanisms underlie a number of fast responses of the human myocardium to mechanical exposures within the TP+M model, this fact does not at all mean that other subcellular mechanisms, too, cannot contribute to these responses. In particular, the stretch-induced modulation of ryanodine receptor activity seems to be one of such additional mechanisms. Fabiato was, probably, the first to argue that calcium-induced calcium release from the sarcoplasmic reticulum may be length dependent [87]. Kohl and Iribe confirmed this fact in experiments on ventricular myocytes of the guinea pig and justified it mathematical modeling [11, 88].

Perhaps, it is just the stretch-induced modulation of ryanodine receptor activity that is responsible for the above-mentioned higher peaks of the Ca^2+^ transients in preloaded twitches compared to isometric conditions which were observed by Vahl et al [55] in the human myocardium preparation from a patient with dilated cardiomyopathy. A similar difference between the Ca^2+^ transient peaks of preloaded and isometric twitches in rat single cardiomyocytes was reported by Yasuda et al [66].

It should be emphasized that the available data on the presence or absence of these differences between Ca^2+^ transient peaks in the myocardium of various animals are quite inconsistent. Let us consider them in a bit more detail. Lab, Allen and Orchard demonstrated the absence of these differences in experiments on cat papillary muscles [66]. White, Boyett and Orchard showed the same for single ventricular myocytes of the guinea-pig [89]. As was mentioned in the foregoing, Lab, Allen and Orchard were the first to report that the shortening of a muscle under low load causes in increase in the free intracellular calcium concentration compared with isometric conditions [58]. That increase, however, occurred in their experiments only some time after the Ca^2+^ transient peak and was present until the end of the muscle shortening phase while the Ca^2+^ transient peaks actually coincided under isometric conditions and during low loaded contractions.

In contrast to these findings, in [66] the increase in calcium compared with the isometric Ca^2+^ transient revealed itself in response to the shortening already during the peak and continued into the early phase of Ca^2+^ transient decay, following which the Ca^2+^ transient curves crossed over.

The data by Vahl et al. for human myocardium preparations demonstrate the presence of considerable differences between the Ca^2+^ transient peaks in some cardiac diseases (dilated cardiomyopathy [55], mitral valve incompetence [65]) and their absence or, in any case, very weak manifestation in others (mitral valve stenosis [65]). Interestingly, in none of these cases do the Ca^2+^ transient curves cross over at the late phase of Ca^2+^ transient decay unlike in the report by [66].

Various suggestions may be made concerning the question why different studies produce so much differing results. For instance, Yasuda et al in their paper proposed separate and quite different hypothetical explanations for the result obtained by Lab et al [58] and for that of White et al [89].

In the first case, they consider the result reported by Lab et al. to be the consequence of papillary muscle non-uniformity, which is likely to lead to a non-uniform distribution of Ca^2+^ indicators and non-uniform mechanical conditions of the component myocytes. Non-uniformity is, no doubt, a very important factor. But why does it lead to the disappearance of only visible differences between the Ca^2+^ transient peaks of isometric and isotonic low-loaded contractions with the differences between the decay phases of the same Ca^2+^ transients in the same muscles remaining clearly visible?

As for why the Ca^2+^ transient peaks matched in the experiments on guinea pig single cardiomyocytes reported by White et al [89], Yasuda and co-authors suggest that the explanation could be associated with certain weaknesses in the experimental setup of White et al.: ªin their isotonic mode (unloaded condition) myocytes were laid on the bottom of the experimental chamber … Such unattached cells often adhere to the surface to create external load». Does it mean that White et al. dealt, in fact, with hidden high-loaded contractions rather than with preloaded or even low-loaded ones? This supposition seems plausible, but just as a supposition rather than an established fact. In other words, the explanations proposed by Yasuda et al. are reasonable but questionable.

These differences could have other explanations, too. For instance, they could be a consequence of species-specificity. Another possible explanation is that the feature of cardiomyocytes being discussed could depend on the region of their localization in the ventricular wall. In discussing the Ca^2+^ transient bumps below, we also consider the experimental study [90], which demonstrated that certain features of Ca^2+^ transients do depend on the region of cardiomyocyte location. However, in relation to the question of Ca^2+^ transient peak match/mismatch in different experimental studies, any such explanations would be purely speculative. What we have today is inconsistent experimental data, and this fact calls for further research. In this context, modeling results cannot fit all experimental data. Our model fits the data presented by Lab et al [58] and White et al [89]. It is valid to say that this is a model limitation, which is unavoidable in this situation. As was already mention above, the introduction of mechano-dependent calcium release from the SR into the model could help realize the opposite variant, i.e. allow for the difference between calcium peaks. In this case, however, they model would not cover their coincidence, a variant we have no reason to neglect. Apparently, additional experimental studies need to be carried out to be able to determine whether a particular context requires one of the alternative variants or an opposite one.

It is interesting to compare the results of our simulation with another of the above-mentioned features of the experimental data from Yasuda et al [66], namely: crossover of Ca^2+^ transient decays in isometric and preloaded isotonic twitches. The lowest afterload in our simulations was 10% of the active isometric force, which is regarded to be already low. However, it was twice higher than the applied preload. In other words, our simulations of afterloaded contractions are not absolutely similar to unloaded isotonic contractions studied by Yasuda et al. For a more detailed comparison with the findings of Yasuda et al., we also reproduced their protocol in the model. Specifically, we simulated a series of contractions in which each odd twitch was isometric and each even twitch was a preloaded isotonic one; besides, both contraction and relaxation took place only under a preload. The results of this numerical experiment are shown in Fig. 13. The right-hand panel in this figure shows Ca^2+^ transients recorded for the last two contractions in the series shown in the left-hand panel. There is a crossover at the end of the fast decay phase of the Ca^2+^ transient corresponding to a preloaded twitch. Qualitatively, this crossover conforms to that presented by Yasuda et al. Of course, there is no complete quantitative match. The divergence of the Ca^2+^ transient traces after the crossover in the paper by Yasuda et al appears to be somewhat greater than in our model.

**Fig. 13.**
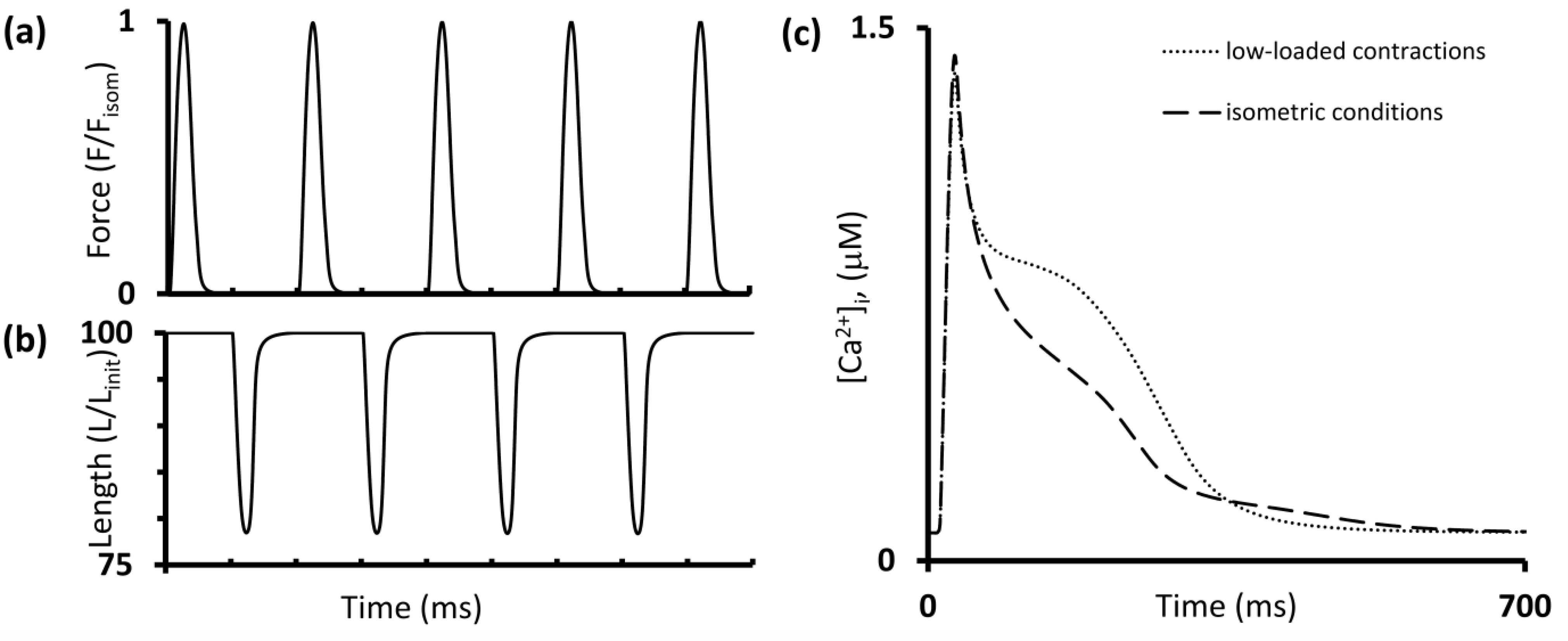
Simulation of different modes of contractions in the TP+M model with the same protocol as applied to rat single cardiac myocytes [66]. According to this protocol, one low-loaded shortening follows one isometric force generation for 90%*L_max_* initial length with the pacing rate of 1 Hz. Stimulation moments are labeled on the time axes. **a** The steady state signals for developed force (as compared to the peak isometric force *F*_*isom*_). **b** The steady state signals for sample length. **c** Ca^2+^ transients for the last isometric twitch (dashed line) and low-loaded shortening (dotted line) are superimposed

As a matter of fact, the recording of calcium fluorescence signals for the preloaded twitch in Fig. 2 in this paper is rather ‘noisy’, which makes difficult its quantitative assessment in the time interval after the crossover. However, we do not claim that the TP+M model can simulate all MCF and MEF effects not only qualitatively but also quantitative to absolute accuracy. On the other hand, we cannot rule it out (although we cannot assert it either) that in is precisely in this particular case the quantitative distinctions from the data of Yasuda et al could be a reflection of species-specificity rather than a consequence of the model’s quantitative imprecision. Our model is oriented towards imitation of human myocardium properties while Yasuda et al. dealt with rat cardiomyocytes.

The latter simulation (Fig. 13) displays a considerable bump on the Ca^2+^ transient during its decay in a preloaded twitch. Similar bumps was obtained in TP+M model for afterloaded twitches. Bumps, though less pronounced, also arose in the simulation оf isometric conditions. The middle panel Fig. 3b shows the numerical experiment result (dashed line in the panel for Ca^2+^ transients) pointing out that within the framework of the models the bump is a consequence of sarcomere shortening: when the model met all isometric conditions for the sarcomeres (i.e. when they were fully fixed during the contraction-relaxation cycle), the bump was gone.

Let us discuss how these modeling results fit experimental data. Previously, such a bump was repeatedly observed in experiments involving myocardium preparations from various species: [76] (Fig. 2), [90–93], [94] (Fig. 5E), [21] (Fig, 12C, experiments). Moreover, the latter two publications present experimental findings obtained on preparations from human hearts. The above data enable us to state confidently that the bump is not an artifact.

At the same time, no bumps are observed in a great number of studies. Of crucial importance for suggesting a possible explanation of the so much diverging experimental data is the study by Choi & Salama [90], who employed optical mapping methods to construct a general map of calcium transients in different regions of the heat. Fig. 2D in this study clearly shows bumps at the base of the guinea pig heart and their absence at the apex. Panel B of the same figure shows in more detail the trace of the Ca^2+^ transient from the central region with a clear-cut bump.

Some of the above studies examined the bump phenomenon specially and point to a range of factors, which are responsible for its extent, including its diminution down to complete disappearance. For instance, Jiang et al [91] examined bumps for temperature-dependence using rat trabeculae and demonstrated that at room temperature the bump was absent from the muscles. One of the most important findings of Jiang et al. was the effect of mechanical conditions of contraction on the extent of the bump on the Ca^2+^ transient [91]. In addition to various experimental protocols in which the preparation loading conditions and length were varied, they evaluated very thoroughly the effect of the mechanics on the extent of the bump using rat ventricular trabeculae treated with various doses of 2,3-butanedione monoxime (BDM). BDM inhibits the mechanical activity of myocardial preparations since it diminishes the number of attached force generating cross bridges. Increasing the concentration of BDM in their experiments reduced active isometric force developed by the trabecula and, at the same time, reduced the extent of the bump (see Fig. 3 in Jiang et al [91]). Finally, at a BDM concentration of 10 mM, the bump vanished at all.

Jiang et al suggested that the mechano-dependence of the bump on the Ca^2+^ transient they observed could be explained by cooperativity between attached cross bridges and Ca^2+^ affinity for TnC [91]. This is in good agreement with our model, in which this cooperativity underlies all manifestations of MCF, the mechanodependence of the bump included.

McDonald et al showed that the extent of the bump on the Ca^2+^ transient could be species-specific [92].Thus, Fig. 8 in their paper shows Ca^2+^ transients measured in isolated cardiomyocytes from porcine and mice hearts. The porcine Ca^2+^ transients in this figure display pronounced bumps: only 2 out of the 14 tested samples did not have bumps, and in the other 12 cells they were pronounced up to producing a secondary increase in calcium in 4 out of the 14 cells. Thus, in the absolute majority of cases, Ca^2+^ transients in the porcine cardiomyocytes revealed significant bumps. In contrast to porcine Ca^2+^ transients, not a single bump arose in any of the 40 tested mice cardiomyocytes.

The current study was focused on fast responses of the myocardium to mechanical impacts and, therefore, we did not deal with slow force responses such as the experimentally shown transient process that occurs during a series of contractions after a rapid increase in length. This process was observed, for example, in experiments on rat and cat myocardium preparations [76, 95]. In essence, the slow force response referred to above is as follows. As a muscle preparation is stretched from a shorter length to a longer one, the peak isometric force increases not only in the first contraction immediately following this muscle stretch but also keeps increasing, not very much but quite noticeably, beat-to-beat in a series of subsequent isometric contractions with a fixed stimulation frequency over a relatively long transient process. Along with force, the [Ca^2+^]_i_ peak also gradually increases beat-to-beat throughout this process.

These slow responses of force and calcium seem impossible to explain by means of the length dependent kinetics of Ca-TnC alone. To reproduce and explain them within the framework of any mathematical model, some additional intracellular mechanisms should therefore be embedded in it. In particular, SACs may mediate Ca^2+^ entry during the slow response. It has been assumed, for example, that a stretch-induced increase in [Ca^2+^]i during this transient process may occur either directly or indirectly by conducting Na^+^ through the SACs, which in turn stimulates the outward NCX current [96]. Not all researchers think that SACs are the main factor underlying the slow response discussed [76, 97]. It could be helpful to verify this hypothesis in the framework of mathematical modeling. However, we are convinced that our model would require a truly complex work to be carried out to do this rather than just fitting the model to the desired result by including a hypothetical stretch-activated current into it. Such a current would a nebulous entity that, in any particular case, would be transfigured and attributed to specific convenient properties *a priori* suitable for obtaining this result.

A complex study of this kind should, undoubtedly, take into account that “despite many experiments confirming the presence of mechanosensitive channels in cardiomyocytes [79, 98–100], there is a large variation in reports of both their characterization and their effects on AP form and duration” [101]. Particularly, “APD has been shown both to increase [98, 102] and decrease [103] in response to the axial strain” [101]. According to this study, the most prevalent SACs are instantly-activating, non-inactivating, cation-selective ones conducting potassium and sodium currents. Referring to various experimental studies, the authors of this paper emphasize that the reversal potentials of stretch-activated currents range from −75 to +10 mV [104], and conductance ranges from 10 to 200 pS [104]. Therefore, for example, even “the inward rectifier K^+^ current has been observed both to increase [98] and decrease [105] in response to different mechanical stimuli” [101]. Thus, the contribution of various SACs to AP development can prove to be both repolarizing and depolarizing, depending on the moment at which the mechanical impact is applied, on its duration and magnitude, and, most importantly, on the magnitude of the reversal potential.

Given this extent of uncertainty and diversity of experimental data, SACs with initially undefined values of the parameter responsible for the reversal potentials and conductance of these channels can be incorporated in a model that has MCF and MEF loops based on the length dependent Ca-TnC kinetics in order to try solving the inverse problem of modeling in relation to these parameters. The latter means finding such parameter values within experimentally determined ranges that would permit one to capture simultaneously the whole range of fast and slow responses of the heart muscle to mechanical impacts. In other words, this means the model should have such values of the parameters for SACs which would allow slow responses to be reproduced without ruining the fast ones in it.

Although the above inverse problem has not yet been solved, the introduction of SACs into various models is still possible and useful. However, this could only be done to simulate relatively local situations pertaining to the stretch since in these cases one can implement only arbitrary values of reversal potentials and conductance (albeit within experimentally reported ranges). For example, it has been shown that SACs thus introduced into a model simulate the slow response of the voltage and [Ca^2+^] to an increase in length [75]. However, this study did not simulate fast responses such as, for example, load dependent relaxation and inactivation of the contractile cycle during quick deformations.

Being reasonable beyond doubt, these approaches leave open the question of the complex role of SACs in the full spectrum of fast and slow responses. Therefore, although the inverse problem formulated above appears to be very difficult to tackle, it is nonetheless important to try and solve it. Indeed, considering the diversity of experimental data available, it could be very useful for the purposes of electrophysiology to obtain evidence-based values for both reversal potentials and conductance for SACs in an integrative study of fast and slow responses by means a mathematical model.

To sum up, this paper has dealt with the question of whether the mechanisms underlying the length dependent cooperative influence of cross-bridge attachment on Ca-TnC kinetics may by themselves produce both MEF and MCF in the fast responses of the human cardiomyocyte to mechanical impacts. In answering it, we believe it was methodologically quite appropriate to include only these mechanisms into the model. This does not mean that the inclusion of SACs into the model will never make sense for simulating fast responses. On the contrary, if in the future we introduce these channels into the TP+M model and try solving the above inverse problem, we will be able to evaluate whether they modulate fast responses, and if so, which ones. On the other hand, it would be imperative to ensure that the model simulates fast responses correctly after incorporating SACs into it and, therefore, it would be essential to perform fundamental testing of these channels for rejecting unacceptable hypothetical properties, which could inevitable initially due to the above-mentioned enormous uncertainty concerning these properties.

## Conclusion

The TP+M model predicts that mechano-calcium, mechano-electric and mechano-mechanical feedbacks are to be inherent in the human cardiomyocyte. The key link underlying these feedbacks observed in the TP+M model during fast responses to mechanical impacts is the cooperativity of regulatory and contractile proteins. These feedbacks arise due to the following mechanisms implemented in the model: cardiomyocyte shortening influences Ca-TnC kinetics and, thus, intracellular calcium kinetics and, via the NCX as a trigger, results in APD length/load dependence.

## Supporting information

Model Description

## List of abbreviation

AP, APs: Action potential, action potentials
APD: Action potential duration
APD_90_: Action potential duration calculated at the level of 90% of repolarization
BDM: 2,3-butanedione monoxime
Ca-TnC: Calcium- troponin C complexes
[Ca^2+^]_i_: Free cytosolic calcium concentration
ECC: Excitation-contraction coupling
F_isom_: Peak isometric force
F_isot_: Level of afterload
i_NaCa_: Na^+^-Ca^2+^ exchanger current
LDI: Load dependence index
L_init_: Initial virtual sample length
L_max_: Length corresponds to a sarcomere length equal to 2.23 μm
MCF: Mechano-calcium feedback
MEF: Mechano-electric feedback
NCX: Na^+^-Ca^2+^exchanger
SACs: Stretch-activated channels
SR: Sarcoplasmic reticulum
SS: Subspace
tCa_70_: Time from peak of Ca^2+^ transient to 70% decay.
TnC: Troponin C
TP: model ‘ten Tusscher – Panfilov’ ionic model
TP+M model: Developed electromechanical model
TTP: Time to peak of twitch
t_50_, t_70_: Time to 50% and 70% force decay from peak force in isometric cycles
Xb, Xb-s: Cross-bridge, cross-bridges

## Author contributions

All authors contributed to conception and design of research, analyzed data, and interpreted the results of numerical experiments; NBV and LB developed the TP+M model; NBV performed numerical experiments; LB and NBV drafted the manuscript; all the authors edited, revised the manuscript, and approved the final manuscript.

## Acknowledgements

Not applicable

## Availability of data and materials

The authors declare that all equations of the developed mathematical model, its parameters and initial values of phasic variables are available within the article and its supplementary information file.

## Compliance with ethical standards

### Ethics approval and consent to participate

Not applicable

### Consent for publication

Not applicable

## Footnotes

4 https://models.physiomeproject.org/exposure/a7179d94365ff0c9c0e6eb7c6a787d3d/ten_tusscher_model_2006_IK1Ko_M_units.cellml/view

## References

1. Orini M, Nanda A, Yates M, Di Salvo C, Roberts N, Lambiase PD, Taggart P (2017) Mechano-electrical feedback in the clinical setting: Current perspectives. Prog Biophys Mol Biol 130:365–375

2. Pfeiffer ER, Tangney JR, Omens JH, McCulloch AD (2014) Biomechanics of cardiac electromechanical coupling and mechanoelectric feedback. J Biomech Eng 136:021007

3. Weckström M, Tavi P, Lab MJ (2007) Mechanoelectric Transduction/Feedback: Physiology and Pathophysiology. In: Weckström M, Tavi P (eds) Cardiac Mechanotransduction. Landes Bioscience and Springer Science+Business Media, pp 48–77

4. Quinn TA, Kohl P, Ravens U (2014) Cardiac mechano-electric coupling research: fifty years of progress and scientific innovation. Prog Biophys Mol Biol 115:71–75

5. Markhasin V, Nikitina L, Routkevich S, Katsnelson L, Schroder E, Keller B (2002) Effects of mechanical interaction between two rabbit cardiac muscles connected in parallel. Gen Physiol Biophys 21:277–302

6. Markhasin VS, Solovyova O, Katsnelson LB, Protsenko Y, Kohl P, Noble D (2003) Mechano-electric interactions in heterogeneous myocardium: development of fundamental experimental and theoretical models. Prog Biophys Mol Biol 82:207–220

7. Protsenko YL et al. (2005) Hybrid duplex: a novel method to study the contractile function of heterogeneous myocardium. Am J Physiol Heart Circ Physiol 289:H2733–2746

8. Solovyova O et al. (2006) Activation sequence as a key factor in spatio-temporal optimization of myocardial function. Philos Trans A Math Phys Eng Sci 364:1367–1383

9. Solovyova O et al. (2003) Mechanical interaction of heterogeneous cardiac muscle segments in silico: effects on Ca2+ handling and action potential. Int J Bifurcat Chaos 13:3757–3782

10. Vikulova NA, Katsnelson LB, Kursanov AG, Solovyova O, Markhasin VS (2016) Mechano-electric feedback in one-dimensional model of myocardium. J Math Biol 73:335–366

11. Iribe G et al. (2009) Axial stretch of rat single ventricular cardiomyocytes causes an acute and transient increase in Ca2+ spark rate. Circ Res 104:787–795

12. Allen D, Kentish J (1988) Calcium concentration in the myoplasm of skinned ferret ventricular muscle following changes in muscle length. J Physiol 407:489–503

13. Gordon AM, Regnier M, Homsher E (2001) Skeletal and cardiac muscle contractile activation: tropomyosin “rocks and rolls”. News Physiol Sci 16:49–55

14. Peyronnet R, Nerbonne JM, Kohl P (2016) Cardiac Mechano-Gated Ion Channels and Arrhythmias. Circ Res 118:311–329

15. Kaufmann R, Lab M, Hennekes R, Krause H (1971) Feedback interaction of mechanical and electrical events in the isolated mammalian ventricular myocardium (cat papillary muscle). Pflügers Archiv 324:100–123

16. Levine JH, Guarnieri T, Kadish AH, White RI, Calkins H, Kan JS (1988) Changes in myocardial repolarization in patients undergoing balloon valvuloplasty for congenital pulmonary stenosis: evidence for contraction-excitation feedback in humans. Circulation 77:70–77

17. Taggart P et al. (1988) Monophasic action potentials at discontinuation of cardiopulmonary bypass: evidence for contraction-excitation feedback in man. Circulation 77:1266–1275

18. Priebe L, Beuckelmann DJ (1998) Simulation study of cellular electric properties in heart failure. Circ Res 82:1206–1223

19. Grandi E, Pasqualini FS, Bers DM (2010) A novel computational model of the human ventricular action potential and Ca transient. J Mol Cell Cardiol 48:112–121

20. ten Tusscher KH, Panfilov AV (2006) Alternans and spiral breakup in a human ventricular tissue model. Am J Physiol Heart Circ Physiol 291:H1088–1100

21. O’Hara T, Virag L, Varro A, Rudy Y (2011) Simulation of the undiseased human cardiac ventricular action potential: model formulation and experimental validation. PLoS Comput Biol 7:e1002061

22. De Oliveira B, Rocha B, Barra L, Toledo E, Sundnes J, Weber dos Santos R (2013) Effects of deformation on transmural dispersion of repolarization using in silico models of human left ventricular wedge. Int J Numer Method Biomed Eng 29:1323–1337

23. Lascano EC, Said M, Vittone L, Mattiazzi A, Mundina-Weilenmann C, Negroni JA (2013) Role of CaMKII in post acidosis arrhythmias: a simulation study using a human myocyte model. J Mol Cell Cardiol 60:172–183

24. Timmermann V, Dejgaard LA, Haugaa KH, Edwards AG, Sundnes J, McCulloch AD, Wall ST (2017) An integrative appraisal of mechano-5electric feedback mechanisms in the heart. Prog Biophys Mol Biol 130:404–417

25. Himeno Y, Asakura K, Cha CY, Memida H, Powell T, Amano A, Noma A (2015) A human ventricular myocyte model with a refined representation of excitation-contraction coupling. Biophys J 109:415–427

26. Rice JJ, Wang F, Bers DM, De Tombe PP (2008) Approximate model of cooperative activation and crossbridge cycling in cardiac muscle using ordinary differential equations. aBiophys J 95:2368–2390

27. Land S, Park-Holohan SJ, Smith NP, Dos Remedios CG, Kentish JC, Niederer SA (2017) A model of cardiac contraction based on novel measurements of tension development in human cardiomyocytes. J Mol Cell Cardiol 106:68–83

28. Sulman T, Katsnelson LB, Solovyova O, Markhasin VS (2008) Mathematical modeling of mechanically modulated rhythm disturbances in homogeneous and heterogeneous myocardium with attenuated activity of Na^+^-K^+^ pump. Bull Math Biol 70:910–949

29. Kazbanov IV, ten Tusscher KH, Panfilov AV (2016) Effects of Heterogeneous Diffuse Fibrosis on Arrhythmia Dynamics and Mechanism. Sci Rep 6:20835

30. Dutta S, Minchole A, Zacur E, Quinn TA, Taggart P, Rodriguez B (2016) Early afterdepolarizations promote transmural reentry in ischemic human ventricles with reduced repolarization reserve. Prog Biophys Mol Biol 120:236–248

31. Perez Alday EA, Whittaker DG, Benson AP, Colman MA (2019) Effects of Heart Rate and Ventricular Wall Thickness on Non-invasive Mapping: An in silico Study. Front Physiol 10:308

32. Vandersickel N, Watanabe M, Tao Q, Fostier J, Zeppenfeld K, Panfilov AV (2018) Dynamical anchoring of distant arrhythmia sources by fibrotic regions via restructuring of the activation pattern. PLoS Comput Biol 14:e1006637

33. Zhan HQ, Xia L, Shou GF, Zang YL, Liu F, Crozier S (2014) Fibroblast proliferation alters cardiac excitation conduction and contraction: a computational study. J Zhejiang Univ Sci B 15:225–242

34. Katsnelson LB, Nikitina LV, Chemla D, Solovyova O, Coirault C, Lecarpentier Y, Markhasin VS (2004) Influence of viscosity on myocardium mechanical activity: a mathematical model. J Theor Biol 230:385–405

35. McDonald KS, Moss RL (1995) Osmotic compression of single cardiac myocytes eliminates the reduction in Ca2+ sensitivity of tension at short sarcomere length. Circ Res 77:199–205

36. Wang YP, Fuchs F (1995) Osmotic compression of skinned cardiac and skeletal muscle bundles: effects on force generation, Ca2+ sensitivity and Ca2+ binding. J Mol Cell Cardiol 27:1235–1244

37. Mateja RD, Greaser ML, de Tombe PP (2013) Impact of titin isoform on length dependent activation and cross-bridge cycling kinetics in rat skeletal muscle. Biochim Biophys Acta 1833:804–811

38. Ait-Mou Y, Hsu K, Farman GP, Kumar M, Greaser ML, Irving TC, de Tombe PP (2016) Titin strain contributes to the Frank-Starling law of the heart by structural rearrangements of both thin- and thick-filament proteins. Proc Natl Acad Sci U S A 113:2306–2311

39. Noble D, Varghese A, Kohl P, Noble P (1998) Improved guinea-pig ventricular cell model incorporating a diadic space, IKr and IKs, and length- and tension-dependent processes. Can J Cardiol 14:123–134

40. Katsnelson LB, Sulman T, Solovyova O, Markhasin VS (2011) Role of myocardial viscoelasticity in disturbances of electrical and mechanical activity in calcium overloaded cardiomyocytes: mathematical modeling. J Theor Biol 272:83–95

41. Izakov V, Katsnelson LB, Blyakhman FA, Markhasin VS, Shklyar TF (1991) Cooperative effects due to calcium binding by troponin and their consequences for contraction and relaxation of cardiac muscle under various conditions of mechanical loading. Circ Res 69:1171–1184

42. Brixius K, Hoischen S, Reuter H, Lasek K, Schwinger RH (2001) Force/shortening-frequency relationship in multicellular muscle strips and single cardiomyocytes of human failing and nonfailing hearts. J Card Fail 7:335–341

43. Gauthier C, Tavernier G, Charpentier F, Langin D, Le Marec H (1996) Functional beta3-adrenoceptor in the human heart. J Clin Invest 98:556–562

44. Gwathmey JK et al. (1991) Diastolic dysfunction in hypertrophic cardiomyopathy. Effect on active force generation during systole. J Clin Invest 87:1023–1031

45. Hasenfuss G, Mulieri LA, Leavitt BJ, Allen PD, Haeberle JR, Alpert NR (1992) Alteration of contractile function and excitation-contraction coupling in dilated cardiomyopathy. Circ Res 70:1225–1232

46. Hasenfuss G, Mulieri LA, Blanchard EM, Holubarsch C, Leavitt BJ, Ittleman F, Alpert NR (1991) Energetics of isometric force development in control and volume-overload human myocardium. Comparison with animal species. Circ Res 68:836–846

47. Holubarsch C et al. (1998) Shortening versus isometric contractions in isolated human failing and non-failing left ventricular myocardium: dependency of external work and force on muscle length, heart rate and inotropic stimulation. Cardiovasc Res 37:46–57

48. Janssen PM, Lehnart SE, Prestle J, Hasenfuss G (1999) Preservation of contractile characteristics of human myocardium in multi-day cell culture. J Mol Cell Cardiol 31:1419–1427

49. Mulieri LA, Hasenfuss G, Leavitt B, Allen PD, Alpert NR (1992) Altered myocardial force-frequency relation in human heart failure. Circulation 85:1743–1750

50. Phillips PJ, Gwathmey JK, Feldman MD, Schoen FJ, Grossman W, Morgan JP (1990) Post-extrasystolic potentiation and the force-frequency relationship: differential augmentation of myocardial contractility in working myocardium from patients with end-stage heart failure. J Mol Cell Cardiol 22:99–110

51. Pieske B et al. (1995) Alterations in intracellular calcium handling associated with the inverse force-frequency relation in human dilated cardiomyopathy. Circulation 92:1169–1178

52. Pieske B et al. (1996) Diminished post-rest potentiation of contractile force in human dilated cardiomyopathy. Functional evidence for alterations in intracellular Ca2+ handling. J Clin Invest 98:764–776

53. Schouten VJ, ter Keurs HE, Quaegebeur JM (1990) Influence of electrogenic Na/Ca exchange on the action potential in human heart muscle. Cardiovasc Res 24:758–767

54. Schwinger RH et al. (1994) The failing human heart is unable to use the Frank-Starling mechanism. Circ Res 74:959–969

55. Vahl CF, Bonz A, Timek T, Hagl S (1994) Intracellular calcium transient of working human myocardium of seven patients transplanted for congestive heart failure. Circ Res 74:952–958

56. Vahl CF et al. (1997) Myocardial length-force relationship in end stage dilated cardiomyopathy and normal human myocardium: analysis of intact and skinned left ventricular trabeculae obtained during 11 heart transplantations. Basic Res Cardiol 92:261–270

57. Milani-Nejad N et al. (2015) The Frank-Starling mechanism involves deceleration of cross-bridge kinetics and is preserved in failing human right ventricular myocardium. Am J Physiol Heart Circ Physiol 309:H2077–2086

58. Lab MJ, Allen DG, Orchard CH (1984) The effects of shortening on myoplasmic calcium concentration and on the action potential in mammalian ventricular muscle. Circ Res 55:825–829

59. Lecarpentier Y, Waldenstrom A, Clergue M, Chemla D, Oliviero P, Martin JL, Swynghedauw B (1987) Major alterations in relaxation during cardiac hypertrophy induced by aortic stenosis in guinea pig. Circ Res 61:107–116

60. Coppini R et al. (2013) Late sodium current inhibition reverses electromechanical dysfunction in human hypertrophic cardiomyopathy. Circulation 127:575–584

61. Gwathmey JK, Slawsky MT, Hajjar RJ, Briggs GM, Morgan JP (1990) Role of intracellular calcium handling in force-interval relationships of human ventricular myocardium. J Clin Invest 85:1599–1613

62. Elnakish MT, Canan BD, Kilic A, Mohler PJ, Janssen PM (2017) Effects of zacopride, a moderate IK1 channel agonist, on triggered arrhythmia and contractility in human ventricular myocardium. Pharmacol Res 115:309–318

63. Lab MJ (1980) Transient depolarisation and action potential alterations following mechanical changes in isolated myocardium. Cardiovasc Res 14:624–637

64. Allen DG, Kentish JC (1985) The cellular basis of the length-tension relation in cardiac muscle. J Mol Cell Cardiol 17:821–840

65. Bonz A, Vahl CF, Hagl S (1997) Contractile behaviour and intracellular calcium during afterloaded contraction in mitral valve disease. Thorac Cardiovasc Surg 45:280–286

66. Yasuda S et al. (2003) Unloaded shortening increases peak of Ca2+ transients but accelerates their decay in rat single cardiac myocytes. Am J Physiol Heart Circ Physiol 285:H470–475

67. Lecarpentier YC, Chuck L, Housmans PR, De Clerck NM, Brutsaert DL (1979) Nature of load dependence of relaxation in cardiac muscle. Am J Physiol Heart Circ Physiol 237:H455–H460

68. Brutsaert DL, De Clerck N, Goethals M, Housmans P (1978) Relaxation of ventricular cardiac muscle. J Physiol 283:469–480

69. Dobrunz LE, Berman MR (1994) Effect of temperature on Ca2+-dependent and mechanical modulators of relaxation in mammalian myocardium. Journal of molecular and cellular cardiology 26:243–250

70. Sipido KR, Volders PG, Vos MA, Verdonck F (2002) Altered Na/Ca exchange activity in cardiac hypertrophy and heart failure: a new target for therapy? Cardiovasc Res 53:782–805

71. Weisser-Thomas J, Piacentino V, 3rd, Gaughan JP, Margulies K, Houser SR (2003) Calcium entry via Na/Ca exchange during the action potential directly contributes to contraction of failing human ventricular myocytes. Cardiovasc Res 57:974–985

72. Solovyova O, Vikulova N, Markhasin VS, Kohl P A novel method for quantifying the contribution of different intracellular mechanisms to mechanically induced changes in action potential characteristics. In: International Workshop on Functional Imaging and Modeling of the Heart, 2003. Springer, pp 8–17

73. Kurihara S, Komukai K (1995) Tension‐dependent changes of the intracellular Ca2+ transients in ferret ventricular muscles. The Journal of Physiology 489:617–625

74. Solovyova O, Vikulova N, Markhasin V, Kohl P (2003) A novel method for quantifying the contribution of different intracellular mechanisms to mechanically induced changes in action potential characteristics. Lecture Notes in Computer Science 2674:8–17

75. Niederer SA, Smith NP (2007) A mathematical model of the slow force response to stretch in rat ventricular myocytes. Biophys J 92:4030–4044

76. Kentish JC, Wrzosek A (1998) Changes in force and cytosolic Ca2+ concentration after length changes in isolated rat ventricular trabeculae. J Physiol 506 (Pt 2):431–444

77. Saeki Y, Kurihara S, Hongo K, Tanaka E (1993) Alterations in intracellular calcium and tension of activated ferret papillary muscle in response to step length changes. J Physiol 463:291–306

78. Kohl P, Day K, Noble D (1998) Cellular mechanisms of cardiac mechano-electric feedback in a mathematical model. Can J Cardiol 14:111–119

79. Zabel M, Koller BS, Sachs F, Franz MR (1996) Stretch-induced voltage changes in the isolated beating heart: importance of the timing of stretch and implications for stretch-activated ion channels. Cardiovascular research 32:120–130

80. Grabarek Z, Grabarek J, Leavis PC, Gergely J (1983) Cooperative binding to the Ca2+-specific sites of troponin C in regulated actin and actomyosin. J Biol Chem 258:14098–14102

81. Razumova MV, Bukatina AE, Campbell KB (2000) Different myofilament nearest-neighbor interactions have distinctive effects on contractile behavior. Biophys J 78:3120–3137

82. Fukuda N, Wu Y, Farman G, Irving TC, Granzier H (2005) Titin-based modulation of active tension and interfilament lattice spacing in skinned rat cardiac muscle. Pflugers Arch 449:449–457

83. Fukuda N, Terui T, Ohtsuki I, Ishiwata S, Kurihara S (2009) Titin and troponin: central players in the frank-starling mechanism of the heart. Curr Cardiol Rev 5:119–124

84. Landesberg A, Sideman S (1994) Mechanical regulation of cardiac muscle by coupling calcium kinetics with cross-bridge cycling: a dynamic model. Am J Physiol 267:H779–795

85. Dupuis LJ, Lumens J, Arts T, Delhaas T (2016) Mechano-chemical Interactions in Cardiac Sarcomere Contraction: A Computational Modeling Study. PLoS Comput Biol 12:e1005126

86. Dupuis LJ, Arts T, Prinzen FW, Delhaas T, Lumens J (2018) Linking cross-bridge cycling kinetics to response to cardiac resynchronization therapy: a multiscale modelling study. Europace 20:iii87–iii93

87. Fabiato A, Fabiato F (1975) Dependence of the contractile activation of skinned cardiac cells on the sarcomere length. Nature 256:54–56

88. Iribe G, Kohl P (2008) Axial stretch enhances sarcoplasmic reticulum Ca2+ leak and cellular Ca2+ reuptake in guinea pig ventricular myocytes: experiments and models. Prog Biophys Mol Biol 97:298–311

89. White E, Boyett MR, Orchard CH (1995) The effects of mechanical loading and changes of length on single guinea-pig ventricular myocytes. J Physiol 482 (Pt 1):93–107

90. Choi BR, Salama G (2000) Simultaneous maps of optical action potentials and calcium transients in guinea-pig hearts: mechanisms underlying concordant alternans. J Physiol 529 Pt 1:171–188

91. Jiang Y, Patterson MF, Morgan DL, Julian FJ (1998) Basis for late rise in fura 2 R signal reporting [Ca2+]i during relaxation in intact rat ventricular trabeculae. Am J Physiol 274:C1273–1282

92. McDonald KS, Hanft LM, Domeier TL, Emter CA (2012) Length and PKA Dependence of Force Generation and Loaded Shortening in Porcine Cardiac Myocytes. Biochem Res Int 2012:371415

93. Lookin O, Protsenko YL (2016) The kinetics of cytosolic calcium in the right ventricular myocardium of guinea pigs and rats. Biophysics 61:119–132

94. Lou Q, Fedorov VV, Glukhov AV, Moazami N, Fast VG, Efimov IR (2011) Transmural heterogeneity and remodeling of ventricular excitation-contraction coupling in human heart failure. Circulation 123:1881–1890

95. Allen DG, Kurihara S (1982) The effects of muscle length on intracellular calcium transients in mammalian cardiac muscle. J Physiol 327:79–94

96. Calaghan SC, White E (1999) The role of calcium in the response of cardiac muscle to stretch. Progress in Biophysics and Molecular Biology 71:59–90

97. Cannell MB (2009) Pulling on the heart strings: a new mechanism within Starling’s law of the heart? Circ Res 104:715–716

98. Isenberg G, Kazanski V, Kondratev D, Gallitelli MF, Kiseleva I, Kamkin A (2003) Differential effects of stretch and compression on membrane currents and [Na+]c in ventricular myocytes. Prog Biophys Mol Biol 82:43–56

99. Guharay F, Sachs F (1984) Stretch-activated single ion channel currents in tissue-cultured embryonic chick skeletal muscle. J Physiol 352:685–701

100. Zeng T, Bett GC, Sachs F (2000) Stretch-activated whole cell currents in adult rat cardiac myocytes. Am J Physiol Heart Circ Physiol 278:H548–557

101. Healy SN, McCulloch AD (2005) An ionic model of stretch-activated and stretch-modulated currents in rabbit ventricular myocytes. Europace 7 Suppl 2:128–134

102. Zhang YH, Youm JB, Sung HK, Lee SH, Ryu SY, Ho WK, Earm YE (2000) Stretch-activated and background non-selective cation channels in rat atrial myocytes. J Physiol 523 Pt 3:607–619

103. Lerman BB, Engelstein ED, Burkhoff D (2001) Mechanoelectrical feedback: role of β-adrenergic receptor activation in mediating load-dependent shortening of ventricular action potential and refractoriness. Circulation 104:486–490

104. Hu H, Sachs F (1997) Stretch-activated ion channels in the heart. J Mol Cell Cardiol 29:1511–1523

105. Tan JH, Liu W, Saint DA (2004) Differential expression of the mechanosensitive potassium channel TREK-1 in epicardial and endocardial myocytes in rat ventricle. Experimental physiology 89:237–242

