## Supplementary material for "Mechano-calcium and mechano-electric feedbacks in the human cardiomyocyte analyzed in a mathematical model": Model Description

### 'ten Tusscher – Panfilov' + Mechanics Model

#### State variables

| | Definition | Initial value | Unit |
|---|---|---|---|
| $V$ | membrane potential | $-84.79$ | mV |
| $d$ | voltage-dependent activation gate ($i_{CaL}$) | $3.575 \cdot 10^{-5}$ | dimensionless |
| $f2$ | fast voltage-dependent inactivation gate ($i_{CaL}$) | 0.9994 | dimensionless |
| $fCass$ | intracellular $Ca^{2+}$ inactivation gate ($i_{CaL}$) | 0.99998 | dimensionless |
| $f$ | slow voltage-dependent inactivation gate ($i_{CaL}$) | 0.9843 | dimensionless |
| $R_{prime}$ | proportion of closed $I_{rel}$ channels | 0.9903 | dimensionless |
| $Ca_{SR}$ | sarcoplasmic reticulum $Ca^{2+}$ concentration | 3.26 | mM |
| $Ca_{ss}$ | subspace $Ca^{2+}$ concentration | 0.00018 | mM |
| $CaTnC$ | $Ca^{2+}$-troponin complexes concentration | 0.00112 | mM |
| $h$ | fast inactivation gate ($i_{Na}$) | 0.7326 | dimensionless |
| $j$ | slow inactivation gate ($i_{Na}$) | 0.7314 | dimensionless |
| $m$ | activation gate ($i_{Na}$) | 0.00188 | dimensionless |
| $Xr1$ | activation gate ($i_{Kr}$) | 0.00023 | dimensionless |
| $Xr2$ | inactivation gate ($i_{Kr}$) | 0.46662 | dimensionless |
| $Xs$ | activation gate ($i_{Ks}$) | 0.00336 | dimensionless |
| $r$ | voltage-dependent activation gate ($i_{to}$) | $2.6 \cdot 10^{-8}$ | dimensionless |
| $s$ | voltage-dependent inactivation gate ($i_{to}$) | 0.999998 | dimensionless |
| $K_i$ | intracellular $K^+$ concentration | 134.42 | mM |
| $Na_i$ | intracellular $Na^+$ concentration | 11.16 | mM |
| $Ca_i$ | intracellular $Ca^{2+}$ concentration | 0.000076 | mM |
| $l_1$ | deformation of $CE$ against its slack length | 0.386 | $\mu$m |
| $l_2$ | deformation of $PE$ against its slack length | 0.386 | $\mu$m |
| $l_3$ | deformation of $XSE$ against its slack length | 0.058 | $\mu$m |
| $v$ | velocity of $CE$ deformation | 0 | $\mu$m/ms |
| $w$ | velocity of $PE$ deformation | 0 | $\mu$m/ms |
| $N$ | cross-bridges concentration | $5.69 \cdot 10^{-6}$ | dimensionless |

#### Constants

| | Definition | Value | Unit |
|---|---|---|---|
| $stim_{amp}$ | amplitude of $i_{stim}$ | 52 | pA/pF |
| $stim_{dur}$ | duration of $i_{stim}$ | 1 | ms |
| $stim_{per}$ | periodicity of $i_{stim}$ | 1000 | ms |
| $stim_{start}$ | start of $i_{stim}$ | 10 | ms |
| $F$ | Faraday constant | 96485.3415 | C/M |
| $R$ | gas constant | 8314.472 | mJ·K$^{-1}$·M$^{-1}$ |
| $T$ | temperature | 310 | K |
| $Cm$ | cell capacitance | 0.185 | $\mu$F |
| $V_c$ | cytoplasmic volume | 0.016404 | mm$^3$ |

Constants (continued)

| | Definition | Value | Unit |
|---|---|---|---|
| $V_{sr}$ | sarcoplasmic reticulum volume | 0.001094 | $mm^3$ |
| $V_{ss}$ | subspace volume | 0.0000547 | $mm^3$ |
| $Ca_o$ | intracellular $Ca^{2+}$ concentration | 2 | mM |
| $K_o$ | extracellular $K^+$ concentration | 5.4 | mM |
| $Na_o$ | extracellular $Na^+$ concentration | 140 | mM |
| $g_{CaL}$ | maximal $i_{CaL}$ conductance | 0.00005 | l/(F· s) |
| $g_{bCa}$ | maximal $i_{bCa}$ conductance | 0.000592 | nS/pF |
| $K_{pCa}$ | $Ca_i$ half-saturation constant of $i_{pCa}$ | 0.0005 | mM |
| $g_{pCa}$ | maximal $i_{pCa}$ conductance | 0.1238 | pA/pF |
| $Buf_{sr}$ | total sarcoplasmic buffer concentration | 10 | mM |
| $Buf_{ss}$ | total subspace buffer concentration | 0.4 | mM |
| $Buf_c$ | total (except $CaTnC$) cytoplasmic buffer concentration | 0.13 | mM |
| $EC$ | $Ca_{SR}$ half-saturation constant of $k_{casr}$ | 1.5 | mM |
| $K_{buf_{sr}}$ | $Ca_{SR}$ half-saturation constant for sarcoplasmic buffer | 0.3 | mM |
| $K_{buf_{ss}}$ | $Ca_{SS}$ half-saturation constant for subspace buffer | 0.00025 | mM |
| $K_{buf_c}$ | $Ca_i$ half-saturation constant for cytoplasmic buffer | 0.00085 | mM |
| $V_{leak}$ | maximal $I_{leak}$ conductance | 0.00036 | $ms^{-1}$ |
| $V_{rel}$ | maximal $I_{rel}$ conductance | 0.1224 | $ms^{-1}$ |
| $V_{xfer}$ | maximal $I_{xfer}$ conductance | 0.00456 | $ms^{-1}$ |
| $K_{up}$ | half-saturation constant of $I_{up}$ | 0.00025 | mM |
| $V_{maxup}$ | maximal $I_{up}$ conductance | 0.00765 | mM/ms |
| $k1_{prime}$ | R to O and R1 to I $I_{rel}$ transition rate | 0.15 | $mM^{-2}\cdot ms^{-1}$ |
| $k2_{prime}$ | O to I and R to R1 $I_{rel}$ transition rate | 0.045 | $mM^{-1}\cdot ms^{-1}$ |
| $k3$ | O to I and R to R1 $I_{rel}$ transition rate | 0.06 | $ms^{-1}$ |
| $k4$ | I to O and R1 to I $I_{rel}$ transition rate | 0.005 | $ms^{-1}$ |
| $max_{sr}$ | maximum value of $k_{casr}$ | 2.5 | dimensionless |
| $min_{sr}$ | minimum value of $k_{casr}$ | 1 | dimensionless |
| $\Pi_{min}$ | parameter of $\Pi_{N_A}$ function | 0.02 | dimensionless |
| $s_c$ | parameter of $N_A(CaTnC, N)$ function | 1.0 | dimensionless |
| $TnC_{tot}$ | total concentration of TnC | 0.07 | mM |
| $k_A$ | cooperativity parameter | 28.0 | $mM^{-1}$ |
| $a_{off}$ | maximum rate constant for $CaTnC$ dissociation | 0.17 | $ms^{-1}$ |
| $a_{on}$ | rate constant for $CaTnC$ association | 35.0 | $mM^{-1}\cdot ms^{-1}$ |
| $g_{Na}$ | maximal $i_{Na}$ conductance | 14.838 | nS/pF |
| $g_{bna}$ | maximal $i_{bNa}$ conductance | 0.00029 | nS/pF |
| $g_{K1}$ | maximal $i_{K1}$ conductance | 5.405 | nS/pF |
| $g_{pK}$ | maximal $i_{pK}$ conductance | 0.0146 | nS/pF |
| $g_{Kr}$ | maximal $i_{Kr}$ conductance | 0.153 | nS/pF |

Constants (continued)

| | Definition | Value | Unit |
|---|---|---|---|
| $P_{kna}$ | relative $i_{Ks}$ permeability to $Na^+$ | 0.03 | dimensionless |
| $g_{Ks}$ | maximal $i_{Ks}$ conductance | 0.392 | nS/pF |
| $g_{to}$ | maximal $i_{to}$ conductance | 0.294 | nS/pF |
| $K_{NaCa}$ | maximal $i_{NaCa}$ | 10000 | pA/pF |
| $K_{sat}$ | saturation factor for $i_{NaCa}$ | 0.1 | dimensionless |
| $Km_{Ca}$ | $Ca_i$ half-saturation constant for $i_{NaCa}$ | 1.38 | mM |
| $Km_{Nai}$ | $Na_i$ half-saturation constant for $i_{NaCa}$ | 87.5 | mM |
| $\alpha$ | factor enhancing outward nature of $i_{NaCa}$ | 1 | dimensionless |
| $\gamma$ | voltage dependence parameter of $i_{NaCa}$ | 0.35 | dimensionless |
| $K_{mNa}$ | $Na_i$ half-saturation constant for $i_{NaK}$ | 40 | mM |
| $K_{mK}$ | $K_o$ half-saturation constant for $i_{NaK}$ | 1 | mM |
| $P_{NaK}$ | maximal $i_{NaK}$ | 2.724 | pA/pF |
| $\lambda$ | scale parameter of $F_{CE}$ | 350.0 | AFU |
| $\alpha_1$ | exponential coefficient of $F_{SE}$ | 14.6 | $\mu$m$^{-1}$ |
| $\beta_1$ | linear coefficient of $F_{SE}$ | 4.2 | AFU |
| $\alpha_2$ | exponential coefficient of $F_{PE}$ | 14.6 | $\mu$m$^{-1}$ |
| $\beta_2$ | linear coefficient of $F_{PE}$ | 0.009 | AFU |
| $\alpha_3$ | exponential coefficient of $F_{XSE}$ | 55.0 | $\mu$m$^{-1}$ |
| $\beta_3$ | linear coefficient of $F_{XSE}$ | 0.11 | AFU |
| $\alpha_{vp_l}$ | exponential coefficient of $F_{VS_1}$ | 16.0 | $\mu$m$^{-1}$ |
| $\alpha_{vp_s}$ | exponential coefficient of $F_{VS_1}$ | 16.0 | $\mu$m$^{-1}$ |
| $\beta_{vp_l}$ | linear coefficient of $F_{VS_1}$ | 0.1 | AFU$\cdot$ ms/$\mu$m |
| $\beta_{vp_s}$ | linear coefficient of $F_{VS_1}$ | 0.1 | AFU$\cdot$ ms/$\mu$m |
| $\alpha_{vs_l}$ | exponential coefficient of $F_{VS_2}$ | 46.0 | $\mu$m$^{-1}$ |
| $\alpha_{vs_s}$ | exponential coefficient of $F_{VS_2}$ | 39.0 | $\mu$m$^{-1}$ |
| $\beta_{vs_l}$ | linear coefficient of $F_{VS_2}$ | 20.0 | AFU$\cdot$ ms/$\mu$m |
| $\beta_{vs_s}$ | linear coefficient of $F_{VS_2}$ | 60.0 | AFU$\cdot$ ms/$\mu$m |
| $v_{max}$ | parameter of $p$ function | 0.0055 | $\mu$m/ms |
| $a$ | parameter of $p$ function | 0.25 | dimensionless |
| $d_h$ | parameter of $P_{star}$ function | 0.5 | dimensionless |
| $\alpha_P$ | parameter of $G_{star}$ function | 4.0 | dimensionless |
| $\alpha_G$ | parameter of $G_{star}$ function | 1.0 | dimensionless |
| $k_\mu$ | parameter of $M_A$ function | 0.6 | dimensionless |
| $\mu$ | parameter of $M_A$ function | 3.3 | dimensionless |
| $g_1$ | parameter of $n_1$ function | 0.6 | $\mu$m$^{-1}$ |
| $g_2$ | parameter of $n_1$ function | 0.52 | dimensionless |
| $n1_A$ | parameter of $n_1$ function | 0.5 | dimensionless |
| $n1_B$ | parameter of $n_1$ function | 55 | $\mu$m |
| $n1_C$ | parameter of $n_1$ function | 1 | dimensionless |
| $n1_Q$ | parameter of $n_1$ function | 0.835 | dimensionless |
| $n1_K$ | parameter of $n_1$ function | 1 | dimensionless |
| $n1_\nu$ | parameter of $n_1$ function | 5 | dimensionless |
| $S_0$ | parameter of $L_{oz}$ function | 1.14 | $\mu$m |
| $S_{055}$ | parameter of $L_{oz}$ function | 0.55 | $\mu$m |
| $S_{046}$ | parameter of $L_{oz}$ function | 0.46 | $\mu$m |

Constants (continued)

| | Definition | Value | Unit |
|---|---|---|---|
| $\kappa_0$ | parameter of $\kappa$ function | 2.1 | dimensionless |
| $\kappa_1$ | parameter of $\kappa$ function | 0.55 | dimensionless |
| $\kappa_2$ | parameter of $\kappa$ function | 0.0 | dimensionless |
| $m_0$ | fraction of strongly attached Xb in steady state isometric conditions | 0.9 | dimensionless |
| $q_1$ | parameter of $q$ function | 0.0173 | $\mathrm{ms}^{-1}$ |
| $q_2$ | parameter of $q$ function | 0.259 | $\mathrm{ms}^{-1}$ |
| $q_3$ | parameter of $q$ function | 0.0173 | $\mathrm{ms}^{-1}$ |
| $q_4$ | parameter of $q$ function | 0.015 | $\mathrm{ms}^{-1}$ |
| $\alpha_Q$ | parameter of $q$ function | 10.0 | dimensionless |
| $\beta_Q$ | parameter of $q$ function | 5.0 | dimensionless |
| $x_{st}$ | parameter of $q$ function | 0.964285 | dimensionless |
| $r0$ | preload | 2.552 | AFU (for $L_{init} = 90\% L_{max}$) |
| $F_{aft}$ | afterload | 6.89 | AFU (for $10\% F_{isom}$) |
| $k_{phys\_rel}$ | parameter of $V_{phys\_rel}$ function | 0.05 | $\mathrm{ms}^{-1}$ |
| $a_{phys\_rel}$ | parameter of $V_{phys\_rel}$ function | *calculated* | $\mu$m |
| $t_{phys\_rel}$ | parameter of $V_{phys\_rel}$ function | *calculated* | ms |
| $per_{phys\_rel}$ | parameter of $V_{phys\_rel}$ function | 230 | ms |

$i_{stim}$, stimulating current.

**Calcium currents**:
$i_{CaL}$, L-type $Ca^{2+}$ current;
$i_{bCa}$, background $Ca^{2+}$ current.

**Calcium translocations**:
$I_{rel}$, $Ca^{2+}$ release from the sarcoplasmic reticulum ($SR$) via ryanodine receptors to the subspace ($SS$);
$I_{xfer}$, $Ca^{2+}$ diffusion from $SS$ to the cytoplasm ($C$);
$I_{leak}$, a small $Ca^{2+}$ leakage from the $SR$ to the cytoplasm;
$I_{up}$, $Ca^{2+}$ pumping from the cytoplasm to the $SR$.
O, open conducting state of $I_{rel}$; R, resting closed state of $I_{rel}$; I, inactivated closed state of $I_{rel}$; R1, resting inactivated closed state of $I_{rel}$.

**Calcium buffers**:
$CaB$, buffering by other than $CaTnC$ intracellular ligands;
$CaTnC$, $Ca^{2+}$-troponin C complexes complexes;
$\Pi_{N_A}$, dependence defining cooperativity of the contractile proteins;
$N_A$, average fraction of the attached cross-bridges per one $CaTnC$ complex;
$CaSRB$, calcium buffering in SR;
$CaSSB$, subspace calcium buffering.

**Sodium currents**:
$i_{Na}$, fast $Na^+$ current;
$i_{bNa}$, background $Na^+$ current.

**Potassium currents**:
$i_{K1}$, inward rectifier $K^+$ current;
$i_{to}$, transient outward current;
$i_{Kr}$, $i_{Ks}$, rapid and slow delayed rectifier current;
$i_{pK}$, plateau $K^+$ current.

**Pumps and exchangers**:
$i_{pCa}$, sarcolemmal $Ca^{2+}$ pump current;
$i_{NaCa}$, $Na^+$–$Ca^{2+}$ exchanger current;
$i_{NaK}$, $Na^+$–$K^+$ pump current.

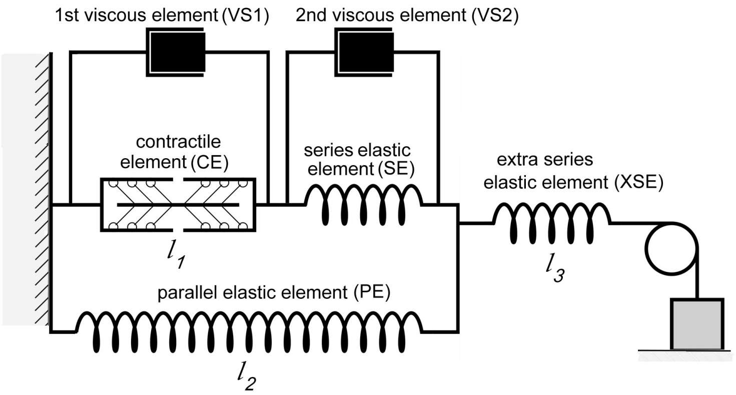


Figure 1: Rheological scheme in the TP+M model

**Forces**: AFU - arbitrary force unit;
$F_{CE}$, contractile element (sarcomere) ($CE$) force;
$F_{SE}$, serial elastic element ($SE$) force;
$F_{PE}$, parallel elastic element ($PE$) force;
$F_{XSE}$, extra serial elastic element ($XSE$) force;
$F_{VS_1}$, $F_{VS_2}$, viscous elements ($VS1$, $VS2$) forces.
$l_1$, deviation of contractile length from its slack length.
$l_2$, deviation of parallel elastic element length from its slack length.
$l_3$, deviation of extra series element length from its slack length.
$l = l_2 + l_3$, deviation of the sample length from its slack length.
$p$, dependence of the average cross-bridge force on the sarcomere shortening/lengthening velocity.
$P_{star}$, dependence of the steady-state sarcomere force on the sarcomere shortening/lengthening velocity.
$G_{star}$, dependence of the steady-state sarcomere stiffness on the velocity.
$M_A$, means end-to-end interaction between adjacent tropomyosin segments in the case if both of them affected by the respective $CaTnC$ complexes formation.
$n_1$, probability of that a myosin head can 'find' a vacant site on the actin filament.
$L_{oz}$, instantaneous length of thick and thin filament overlap zone.
$\kappa$, function required for variation the ratio between rates of cross-bridge attachment and detachment.

$q$, stationary relation 'stiffness-velocity' for the sample.
$L_{init}$, initial length of the sample.
$L_{max}$, corresponds to a sarcomere length equal to 2.23 $\mu m$.
$F_{isom}$, maximum of isometric force at given $L_{init}$.
$V_{phys\_rel}$, dependence of sample length return during physiological relaxation.
$k_{phys\_rel}$, physiological relaxation velocity.
$a_{phys\_rel}$, physiological relaxation amplitude calculated as a difference between end-systolic and initial lengths of the sample.
$t_{phys\_rel}$, time to start physiological relaxation fixed at the moment when $F_{sample} = r0$.
$per_{phys\_rel}$ period of physiological relaxation.

### MODEL EQUATIONS

TNNP BLOCK (with modifications)

MEMBRANE POTENTIAL

$$i_{Stim} = \begin{cases} -stim_{amp} & \text{if } \left(time - \lfloor \frac{time}{stim_{per}} \rfloor \cdot stim_{per} \geq stim_{start}\right) \text{ and} \\ & \text{and } \left(time - \lfloor \frac{time}{stim_{per}} \rfloor \cdot stim_{per} \leq stim_{start} + stim_{dur}\right) \\ 0 & \text{otherwise} \end{cases}$$

$$\frac{\mathrm{d}V}{\mathrm{d}time} = \frac{-1}{1} \cdot (i_{K1} + i_{to} + i_{Kr} + i_{Ks} + i_{CaL} + i_{NaK} + i_{Na} + i_{b_{Na}} + i_{NaCa} + i_{b_{Ca}} + i_{p_K} + i_{p_{Ca}} + i_{Stim})$$

REVERSAL POTENTIALS

$$E_{Na} = \frac{R \cdot T}{F} \cdot \ln \frac{Na_o}{Na_i}$$

$$E_K = \frac{R \cdot T}{F} \cdot \ln \frac{K_o}{K_i}$$

$$E_{Ks} = \frac{R \cdot T}{F} \cdot \ln \frac{K_o + P_{kna} \cdot Na_o}{K_i + P_{kna} \cdot Na_i}$$

$$E_{Ca} = \frac{0.5 \cdot R \cdot T}{F} \cdot \ln \frac{Ca_o}{Ca_i}$$

#### L-type $Ca^{2+}$ current

$$i_{CaL} = g_{CaL} \cdot d \cdot f \cdot f_2 \cdot f_{Cass} \cdot 4 \cdot \frac{(V-15) \cdot F^2}{R \cdot T} \cdot \frac{\left(0.25 \cdot Ca_{ss} \cdot e^{\frac{2 \cdot (V-15) \cdot F}{R \cdot T}} - Ca_o\right)}{e^{\frac{2 \cdot (V-15) \cdot F}{R \cdot T}} - 1}$$

#### L-type $Ca^{2+}$ current. $d$ gate

$$d_{inf} = \frac{1}{1 + e^{\frac{-8-V}{7.5}}}$$

$$\alpha_d = \frac{1.4}{1 + e^{\frac{-35-V}{13}}} + 0.25$$

$$\beta_d = \frac{1.4}{1 + e^{\frac{V+5}{5}}}$$

$$\gamma_d = \frac{1}{1 + e^{\frac{50-V}{20}}}$$

$$\tau_d = 1 \cdot \alpha_d \cdot \beta_d + \gamma_d$$

$$\frac{\mathrm{d}d}{\mathrm{d}time} = \frac{d_{inf} - d}{\tau_d}$$

#### L-type $Ca^{2+}$ current. $f2$ gate

$$f2_{inf} = \frac{0.67}{1 + e^{\frac{V+35}{7}}} + 0.33$$

$$\tau_{f2} = 562 \cdot e^{\frac{-(V+27)^2}{240}} + \frac{31}{1 + e^{\frac{25-V}{10}}} + \frac{80}{1 + e^{\frac{V+30}{10}}}$$

$$\frac{\mathrm{d}f_2}{\mathrm{d}time} = \frac{f2_{inf} - f2}{\tau_{f2}}$$

#### L-type $Ca^{2+}$ current. $fCa_{SS}$ gate

$$fCass_{inf} = \frac{0.6}{1 + \left(\frac{Ca_{ss}}{0.05}\right)^2} + 0.4$$

$$\tau_{fCass} = \frac{80}{1 + \left(\frac{Ca_{ss}}{0.05}\right)^2} + 2$$

### L-TYPE $Ca^{2+}$ CURRENT. $fCa_{SS}$ GATE (CONTINUED)

$$\frac{\mathrm{d}f_{Cass}}{\mathrm{d}time} = \frac{fCass_{inf} - fCass}{\tau_{fCass}}$$

### L-TYPE $Ca^{2+}$ CURRENT. $f$ GATE

$$f_{inf} = \frac{1}{1 + e^{\frac{V+20}{7}}}$$

$$\tau_f = 1102.5 \cdot e^{\frac{-(V+27)^2}{225}} + \frac{200}{1 + e^{\frac{13-V}{10}}} + \frac{180}{1 + e^{\frac{V+30}{10}}} + 20$$

$$\frac{\mathrm{d}f}{\mathrm{d}time} = \frac{f_{inf} - f}{\tau_f}$$

### $Ca^{2+}$ BACKGROUND CURRENT

$$i_{b_{Ca}} = g_{bca} \cdot (V - E_{Ca})$$

### $Ca^{2+}$ PUMP CURRENT

$$i_{p_{Ca}} = \frac{g_{pCa} \cdot Ca_i}{Ca_i + K_{pCa}}$$

### $Ca^{2+}$ DYNAMICS

$$i_{rel} = V_{rel} \cdot O \cdot (Ca_{SR} - Ca_{ss})$$

$$i_{up} = \frac{Vmax_{up}}{1 + \frac{K_{up}^2}{Ca_i^2}}$$

$$i_{leak} = V_{leak} \cdot (Ca_{SR} - Ca_i)$$

$$i_{xfer} = V_{xfer} \cdot (Ca_{ss} - Ca_i)$$

$$O = \frac{k1 \cdot Ca_{ss}^2 \cdot R_{prime}}{k3 + k1 \cdot Ca_{ss}^2}$$

$$\frac{\mathrm{d}R_{prime}}{\mathrm{d}time} = -k2 \cdot Ca_{ss} \cdot R_{prime} + k4 \cdot (1 - R_{prime})$$

$$k1 = \frac{k1_{prime}}{kcasr}$$

$$k2 = k2_{prime} \cdot kcasr$$

$$kcasr = max_{sr} - \frac{max_{sr} - min_{sr}}{1 + \left(\frac{EC}{Ca_{SR}}\right)^2}$$

$$B_{Cabufc} = \frac{1}{1 + \frac{Buf_c \cdot K_{buf_c}}{(Ca_i + K_{buf_c})^2}}$$

$$B_{Cabufsr} = \frac{1}{1 + \frac{Buf_{sr} \cdot K_{buf_{sr}}}{(Ca_{SR} + K_{buf_{sr}})^2}}$$

$$B_{Cabufss} = \frac{1}{1 + \frac{Buf_{ss} \cdot K_{buf_{ss}}}{(Ca_{ss} + K_{buf_{ss}})^2}}$$

$$N_A = \frac{TnC_{tot} \cdot N \cdot sc}{L_{oz} \cdot CaTnC}$$

$$\Pi_{N_A} = \begin{cases} 1 & \text{if } N_A \leq 0 \\ \Pi_{min}^{N_A} & \text{if } N_A \leq 1 \\ \Pi_{min} & \text{otherwise} \end{cases}$$

$$\frac{\mathrm{d}CaTnC}{\mathrm{d}time} = a_{on} \cdot (TnC_{tot} - CaTnC) \cdot Ca_i - a_{off} \cdot e^{-k_A \cdot CaTnC} \cdot \Pi_{N_A} \cdot CaTnC$$

$$\frac{\mathrm{d}Ca_{sr}}{\mathrm{d}time} = B_{Cabufsr} \cdot (i_{up} - (i_{rel} + i_{leak}))$$

$$\frac{\mathrm{d}Ca_{ss}}{\mathrm{d}time} = B_{Cabufss} \cdot \left(\frac{-1 \cdot i_{CaL} \cdot Cm}{2 \cdot 1 \cdot V_{ss} \cdot F} + \frac{i_{rel} \cdot V_{sr}}{V_{ss}} - \frac{i_{xfer} \cdot V_c}{V_{ss}}\right)$$

$$CaB = \frac{Buf_c \cdot Ca_i}{Ca_i + K_{buf_c}}$$

$$CaSRB = \frac{Buf_{sr} \cdot Ca_{sr}}{Ca_{sr} + K_{buf_{sr}}}$$

$$CaSSB = \frac{Buf_{ss} \cdot Ca_{ss}}{Ca_{ss} + K_{buf_{ss}}}$$

#### FAST $Na^+$ CURRENT

$$i_{Na} = g_{Na} \cdot m^3 \cdot h \cdot j \cdot (V - E_{Na})$$

#### FAST $Na^+$ CURRENT. $h$ GATE

$$h_{inf} = \frac{1}{\left(1 + e^{\frac{V+71.55}{7.43}}\right)^2}$$

$$\alpha_h = \begin{cases} 0.057 \cdot e^{\frac{-(V+80)}{6.8}} & \text{if } V < -40 \\ 0 & \text{otherwise} \end{cases}$$

$$\beta_h = \begin{cases} 2.7 \cdot e^{0.079 \cdot V} + 310000 \cdot e^{0.3485 \cdot V} & \text{if } V < -40 \\ \dfrac{0.77}{0.13 \cdot \left(1 + e^{\frac{V+10.66}{-11.1}}\right)} & \text{otherwise} \end{cases}$$

$$\tau_h = \frac{1}{\alpha_h + \beta_h}$$

$$\frac{\mathrm{d}h}{\mathrm{d}time} = \frac{h_{inf} - h}{\tau_h}$$

#### FAST $Na^+$ CURRENT. $j$ GATE

$$j_{inf} = \frac{1}{\left(1 + e^{\frac{V+71.55}{7.43}}\right)^2}$$

$$\alpha_j = \begin{cases} \dfrac{\left(-25428 \cdot e^{0.2444 \cdot V} - 6.948 \cdot 10^{-6} \cdot e^{-0.04391 \cdot V}\right) \cdot (V + 37.78)}{1 + e^{0.311 \cdot (V+79.23)}} & \text{if } V < -40 \\ 0 & \text{otherwise} \end{cases}$$

$$\beta_j = \begin{cases} \dfrac{0.02424 \cdot e^{-0.01052 \cdot V}}{1 + e^{-0.1378 \cdot (V+40.14)}} & \text{if } V < -40 \\ \dfrac{0.6 \cdot e^{0.057 \cdot V}}{1 + e^{-0.1 \cdot (V+32)}} & \text{otherwise} \end{cases}$$

$$\tau_j = \frac{1}{\alpha_j + \beta_j}$$

$$\frac{\mathrm{d}j}{\mathrm{d}time} = \frac{j_{inf} - j}{\tau_j}$$

#### FAST $Na^+$ CURRENT. $m$ GATE

$$m_{inf} = \frac{1}{\left(1 + e^{\frac{-56.86 - V}{9.03}}\right)^2}$$

$$\alpha_m = \frac{1}{1 + e^{\frac{-60 - V}{5}}}$$

$$\beta_m = \frac{0.1}{1 + e^{\frac{V + 35}{5}}} + \frac{0.1}{1 + e^{\frac{V - 50}{200}}}$$

$$\tau_m = 1 \cdot \alpha_m \cdot \beta_m$$

$$\frac{\mathrm{d}m}{\mathrm{d}time} = \frac{m_{inf} - m}{\tau_m}$$

#### $Na^+$ BACKGROUND CURRENT

$$i_{b_{Na}} = g_{bna} \cdot (V - E_{Na})$$

#### INWARD RECTIFIER $K^+$ CURRENT

$$\alpha_{K1} = \frac{0.1}{1 + e^{0.06 \cdot (V - E_K - 200)}}$$

$$\beta_{K1} = \frac{3 \cdot e^{0.0002 \cdot (V - E_K + 100)} + e^{0.1 \cdot (V - E_K - 10)}}{1 + e^{-0.5 \cdot (V - E_K)}}$$

$$xK1_{inf} = \frac{\alpha_{K1}}{\alpha_{K1} + \beta_{K1}}$$

$$i_{K1} = g_{K1} \cdot xK1_{inf} \cdot \sqrt{\frac{K_o}{5.4}} \cdot (V - E_K)$$

#### $K^+$ PLATEAU CURRENT

$$i_{p_K} = \frac{g_{pK} \cdot (V - E_K)}{1 + e^{\frac{25 - V}{5.98}}}$$

#### RAPID TIME DEPENDENT $K^+$ CURRENT

$$i_{Kr} = g_{Kr} \cdot \sqrt{\frac{K_o}{5.4}} \cdot Xr1 \cdot Xr2 \cdot (V - E_K)$$

#### rapid time dependent $K^+$ current. Xr1 gate

$$xr1_{inf} = \frac{1}{1 + e^{\frac{-26-V}{7}}}$$

$$\alpha_{xr1} = \frac{450}{1 + e^{\frac{-45-V}{10}}}$$

$$\beta_{xr1} = \frac{6}{1 + e^{\frac{V+30}{11.5}}}$$

$$\tau_{xr1} = 1 \cdot \alpha_{xr1} \cdot \beta_{xr1}$$

$$\frac{\mathrm{d}Xr1}{\mathrm{d}time} = \frac{xr1_{inf} - Xr1}{\tau_{xr1}}$$

#### rapid time dependent $K^+$ current. Xr2 gate

$$xr2_{inf} = \frac{1}{1 + e^{\frac{V+88}{24}}}$$

$$\alpha_{xr2} = \frac{3}{1 + e^{\frac{-60-V}{20}}}$$

$$\beta_{xr2} = \frac{1.12}{1 + e^{\frac{V-60}{20}}}$$

$$\tau_{xr2} = 1 \cdot \alpha_{xr2} \cdot \beta_{xr2}$$

$$\frac{\mathrm{d}Xr2}{\mathrm{d}time} = \frac{xr2_{inf} - Xr2}{\tau_{xr2}}$$

#### slow time dependent $K^+$ current

$$i_{Ks} = g_{Ks} \cdot Xs^2 \cdot (V - E_{Ks})$$

#### slow time dependent $K^+$ current. Xs gate

$$xs_{inf} = \frac{1}{1 + e^{\frac{-5-V}{14}}}$$

$$\alpha_{xs} = \frac{1400}{\sqrt{1 + e^{\frac{5-V}{6}}}}$$

### slow time dependent $K^+$ current. Xs gate (continued)

$$\beta_{xs} = \frac{1}{1 + e^{\frac{V-35}{15}}}$$

$$\tau_{xs} = 1 \cdot \alpha_{xs} \cdot \beta_{xs} + 80$$

$$\frac{\mathrm{d}Xs}{\mathrm{d}time} = \frac{xs_{inf} - Xs}{\tau_{xs}}$$

### transient outward $K^+$ current

$$i_{to} = g_{to} \cdot r \cdot s \cdot (V - E_K)$$

### transient outward $K^+$ current. r gate

$$r_{inf} = \frac{1}{1 + e^{\frac{20-V}{6}}}$$

$$\tau_r = 9.5 \cdot e^{\frac{-(V+40)^2}{1800}} + 0.8$$

$$\frac{\mathrm{d}r}{\mathrm{d}time} = \frac{r_{inf} - r}{\tau_r}$$

### transient outward $K^+$ current. s gate

$$s_{inf} = \frac{1}{1 + e^{\frac{V+20}{5}}}$$

$$\tau_s = 85 \cdot e^{\frac{-(V+45)^2}{320}} + \frac{5}{1 + e^{\frac{V-20}{5}}} + 3$$

$$\frac{\mathrm{d}s}{\mathrm{d}time} = \frac{s_{inf} - s}{\tau_s}$$

### $Na^+$-$Ca^{2+}$ exchanger current

$$i_{NaCa} = \frac{K_{NaCa} \cdot \left(e^{\frac{\gamma \cdot V \cdot F}{R \cdot T}} \cdot Na_i^3 \cdot Ca_o - e^{\frac{(\gamma-1) \cdot V \cdot F}{R \cdot T}} \cdot Na_o^3 \cdot Ca_i \cdot \alpha\right)}{\left(Km_{Nai}^3 + Na_o^3\right) \cdot (Km_{Ca} + Ca_o) \cdot \left(1 + K_{sat} \cdot e^{\frac{(\gamma-1) \cdot V \cdot F}{R \cdot T}}\right)}$$

### $Na^{+}$-$K^{+}$ PUMP CURRENT

$$i_{NaK} = P_{NaK} \cdot \frac{K_o \cdot Na_i}{(K_o + K_{mk}) \cdot (Na_i + K_{mNa}) \cdot (1 + 0.1245 \cdot e^{\frac{-0.1 \cdot V \cdot F}{R \cdot T}} + 0.0353 \cdot e^{\frac{-V \cdot F}{R \cdot T}})}$$

### $Ca^{2+}$ DYNAMICS

$$\frac{\mathrm{d}Ca_i}{\mathrm{d}t} = B_{Cabufc} \cdot \left( \frac{(i_{leak} - i_{up}) \cdot V_{sr}}{V_c} + i_{xfer} - \frac{(i_{b_{Ca}} + i_{p_{Ca}} - 2 \cdot i_{NaCa}) \cdot Cm}{V_c \cdot F} - \frac{\mathrm{d}CaTnC}{\mathrm{d}t} \right)$$

### $K^{+}$ DYNAMICS

$$\frac{\mathrm{d}K_i}{\mathrm{d}time} = \frac{-1 \cdot (i_{K1} + i_{to} + i_{Kr} + i_{Ks} + i_{pK} + i_{Stim} - 2 \cdot i_{NaK})}{1 \cdot V_c \cdot F} \cdot Cm$$

### $Na^{+}$ DYNAMICS

$$\frac{\mathrm{d}Na_i}{\mathrm{d}time} = \frac{-1 \cdot (i_{Na} + i_{b_{Na}} + 3 \cdot i_{NaK} + 3 \cdot i_{NaCa})}{1 \cdot V_c \cdot F} \cdot Cm$$

### MECHANICAL BLOCK

FORCE

$$F_{CE} = \lambda \cdot p_v \cdot N$$

$$F_{SE} = \beta_1 \cdot \left(e^{\alpha_1 \cdot (l_2 - l_1)} - 1\right)$$

$$F_{PE} = \beta_2 \cdot \left(e^{\alpha_2 \cdot l_2} - 1\right)$$

$$F_{XSE} = \beta_3 \cdot \left(e^{\alpha_3 \cdot l_3} - 1\right)$$

$$F_{VS_1} = k_{P_{vis}} \cdot v$$

$$F_{VS_2} = k_{S_{vis}} \cdot (w - v)$$

$$F_{sample} = F_{XSE}$$

CONTRACTION MODES

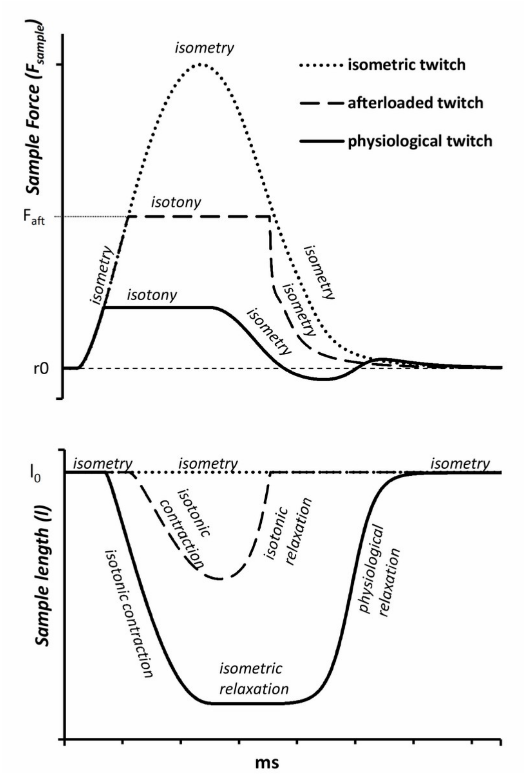


Figure 2: Contraction modes in the TP+M model

$$contraction_{mode} = \begin{cases} isometry & \\ isotony & \text{if } (F_{sample} > F_{aft}) \text{ and } \left(l \le l_0 \cdot \left(1 + 1 \cdot 10^{-4}\right)\right) \\ isometric\_relaxation & \text{if } (l\ is\ end - systolic\ length) \text{ and } (F_{sample} > r0) \\ physiological\_relaxation & \text{starts after isometric relaxation when } (F_{sample} \le r0) \end{cases}$$

LENGTH

$$l = l_2 + l_3$$

$$\frac{\mathrm{d}l_1}{\mathrm{d}time} = v$$

$$\frac{\mathrm{d}l_2}{\mathrm{d}time} = w$$

$$V_{phys\_rel} = \frac{a_{phys\_rel} \cdot k_{phys\_rel} \cdot e^{(-k_{phys\_rel} \cdot (t - (t_{phys\_rel} - per_{phys\_rel}/2)))}}{(1 + e^{(-k_{phys\_rel} \cdot (t - (t_{phys\_rel} - per_{phys\_rel}/2))})^2}$$

$$\frac{\mathrm{d}l_3}{\mathrm{d}time} = \begin{cases} -w & \text{if } contraction_{mode} = isometry\ and\ isometric\_relaxation \\ 0 & \text{if } contraction_{mode} = isotony \\ V_{phys\_rel} - w & \text{if } contraction_{mode} = physiological\_relaxation \end{cases}$$

$$alp_p = \begin{cases} \alpha_{vp_l} & \text{if } v \leq 0 \\ \alpha_{vp_s} & \text{otherwise} \end{cases}$$

$$k_{P_{vis}} = \begin{cases} \beta_{vp_l} \cdot e^{\alpha_{vp_l} \cdot l_1} & \text{if } v \leq 0 \\ \beta_{vp_s} \cdot e^{\alpha_{vp_s} \cdot l_1} & \text{otherwise} \end{cases}$$

$$\phi_\chi = \begin{cases} \frac{-\left(\lambda \cdot K_\kappa \cdot p_v + alp_p \cdot k_{P_{vis}} \cdot v^2 + \left(\alpha_2 \cdot \beta_2 \cdot e^{\alpha_2 \cdot l_2} + \alpha_3 \cdot \beta_3 \cdot e^{\alpha_3 \cdot l_3}\right) \cdot w\right)}{\lambda \cdot N \cdot p_{prime_v} + k_{P_{vis}}}, & \text{if } contraction_{mode} = isometry\ and\ isometric\_relaxation \\ \frac{-\left(\lambda \cdot K_\kappa \cdot p_v + alp_p \cdot k_{P_{vis}} \cdot v^2 + \alpha_2 \cdot \beta_2 \cdot e^{\alpha_2 \cdot l_2} \cdot w\right)}{\lambda \cdot N \cdot p_{prime_v} + k_{P_{vis}}}, & \text{if } contraction_{mode} = isotonic \\ \frac{-\left(\lambda \cdot K_\kappa \cdot p_v + alp_p \cdot k_{P_{vis}} \cdot v^2 + \alpha_2 \cdot \beta_2 \cdot e^{\alpha_2 \cdot l_2} \cdot w - \alpha_3 \cdot \beta_3 \cdot e^{\alpha_3 \cdot l_3} \cdot \left(V_{phys\_rel} - w\right)\right)}{\lambda \cdot N \cdot p_{prime_v} + k_{P_{vis}}}, & \text{if } contraction_{mode} = physiological\_relaxation \end{cases}$$

$$\frac{\mathrm{d}v}{\mathrm{d}time} = \phi_\chi$$

$$alp_s = \begin{cases} \alpha_{vs_l} & \text{if } w \leq v \\ \alpha_{vs_s} & \text{otherwise} \end{cases}$$

$$k_{S_{vis}} = \begin{cases} \beta_{vs_l} \cdot e^{\alpha_{vs_l} \cdot (l_2 - l_1)} & \text{if } w \leq v \\ \beta_{vs_s} \cdot e^{\alpha_{vs_s} \cdot (l_2 - l_1)} & \text{otherwise} \end{cases}$$

$$\frac{\mathrm{d}w}{\mathrm{d}time} = \begin{cases} \phi_\chi - alp_s \cdot (w - v)^2 - \dfrac{\alpha_1 \cdot \beta_1 \cdot e^{\alpha_1 \cdot (l_2 - l_1)} \cdot (w - v) + \left(\alpha_2 \cdot \beta_2 \cdot e^{\alpha_2 \cdot l_2} + \alpha_3 \cdot \beta_3 \cdot e^{\alpha_3 \cdot l_3}\right) \cdot w}{k_{S_{vis}}}, \\ \text{if } (contraction_{mode} = isometry\ and\ isometric\_relaxation) \\ \\ \dfrac{k_{S_{vis}} \cdot \left(\phi_\chi - alp_s \cdot (w - v)^2\right) - \alpha_1 \cdot \beta_1 \cdot e^{\alpha_1 \cdot (l_2 - l_1)} \cdot (w - v) - \alpha_2 \cdot \beta_2 \cdot e^{\alpha_2 \cdot l_2} \cdot w}{k_{S_{vis}}}, \\ \text{if } (contraction_{mode} = isotony) \\ \\ \phi_\chi - alp_s \cdot (w - v)^2 - \\ - \dfrac{\alpha_1 \cdot \beta_1 \cdot e^{\alpha_1 \cdot (l_2 - l_1)} \cdot (w - v) + \alpha_2 \cdot \beta_2 \cdot e^{\alpha_2 \cdot l_2} \cdot w - \alpha_3 \cdot \beta_3 \cdot e^{\alpha_3 \cdot l_3} \cdot (V_{phys\_rel} - w)}{k_{S_{vis}}}, \\ \text{if } (contraction_{mode} = physiological\_relaxation) \end{cases}$$

$$v_1 = \frac{v_{max}}{10}$$

$$\gamma_2 = \frac{a \cdot d_h \cdot \left(\dfrac{v_1}{v_{max}}\right)^2}{3 \cdot a \cdot d_h - \dfrac{(a+1) \cdot v_1}{v_{max}}}$$

$$P_{star} = \begin{cases} \dfrac{a \cdot \left(1 + \dfrac{v}{v_{max}}\right)}{a - \dfrac{v}{v_{max}}} & \text{if } v \leq 0 \\ 1 + d_h - \dfrac{d_h^2 \cdot a}{\dfrac{a \cdot d_h}{\gamma_2} \cdot \left(\dfrac{v}{v_{max}}\right)^2 + \dfrac{(a+1) \cdot v}{v_{max}} + a \cdot d_h} & \text{otherwise} \end{cases}$$

$$G_{star} = \begin{cases} 1 + \frac{0.6 \cdot v}{v_{max}} & \text{if } (v \leq 0) \\ \dfrac{P_{star}}{\dfrac{(0.4 \cdot a + 1) \cdot v}{a \cdot v_{max}} + 1} & \text{if } (0 < v) \text{ and } (v \leq v_1) \\ \dfrac{P_{star} \cdot e^{-\alpha_G \cdot \left(\frac{v - v_1}{v_{max}}\right)^{\alpha_P}}}{\dfrac{(0.4 \cdot a + 1) \cdot v}{a \cdot v_{max}} + 1} & \text{otherwise} \end{cases}$$

$$case_1 = \frac{a \cdot (0.4 + 0.4 \cdot a)}{v_{max} \cdot ((a+1) \cdot 0.4)^2}$$

$$case_2 = \frac{a \cdot 1 \cdot \left(1 + 0.4 \cdot a + \dfrac{1.2 \cdot v}{v_{max}} + 0.6 \cdot \left(\dfrac{v}{v_{max}}\right)^2\right)}{v_{max} \cdot \left(\left(a - \dfrac{v}{v_{max}}\right) \cdot \left(1 + \dfrac{0.6 \cdot v}{v_{max}}\right)\right)^2}$$

$$case_3 = \frac{0.4 \cdot a + 1}{a \cdot v_{max}}$$

$$case_4 = \frac{1}{v_{max}} \cdot e^{-\alpha_G \cdot \left(\frac{v - v_1}{v_{max}}\right)^{\alpha_P}} \cdot \left(\frac{0.4 \cdot a + 1}{a} + \alpha_G \cdot \alpha_P \cdot \left(1 + \frac{(0.4 \cdot a + 1) \cdot v}{a \cdot v_{max}}\right) \cdot \left(\frac{v - v_1}{v_{max}}\right)^{\alpha_P - 1}\right)$$

$$p_{prime_v} = \begin{cases} case_1 & \text{if } v \leq -v_{max} \\ case_2 & \text{if } (-v_{max} < v) \text{ and } (v \leq 0) \\ case_3 & \text{if } (0 < v) \text{ and } (v \leq v_1) \\ case_4 & \text{otherwise} \end{cases}$$

$$p_v = \frac{P_{star}}{G_{star}}$$

$$M_A = \frac{\left(\frac{CaTnC}{TnC_{tot}}\right)^{\mu} \cdot \left(1 + k_{\mu}^{\mu}\right)}{\left(\frac{CaTnC}{TnC_{tot}}\right)^{\mu} + k_{\mu}^{\mu}}$$

$$temp_{n1} = (g_1 \cdot l_1 + g_2) \cdot \left(n1_A + \frac{n1_K - n1_A}{\left(n1_C + n1_Q \cdot e^{-n1_B \cdot l_1}\right)^{\frac{1}{n1_{\nu}}}}\right)$$

$$n_1 = \begin{cases} 0 & \text{if } temp_{n1} < 0 \\ temp_{n1} & \text{if } temp_{n1} < 1 \\ 1 & \text{otherwise} \end{cases}$$

$$L_{oz} = \begin{cases} \frac{l_1 + S_0}{S_{046} + S_0} & \text{if } l_1 \leq S_{055} \\ \frac{S_0 + S_{055}}{S_{046} + S_0} & \text{otherwise} \end{cases}$$

$$\kappa = \begin{cases} \kappa_1 + \kappa_2 \cdot \frac{v}{v_{max}} & \text{if } v < 0 \\ \kappa_1 & \text{otherwise} \end{cases}$$

$$v_{st} = x_{st} \cdot v_{max}$$

$$q_v = \begin{cases} q_1 - \frac{q_2 \cdot v}{v_{max}} & \text{if } v \leq 0 \\ \frac{(q_4 - q_3) \cdot v}{v_{st}} + q_3 & \text{if } (v \leq v_{st}) \text{ and } (0 < v) \\ \frac{q_4}{\left(1 + \frac{\beta_Q \cdot (v - v_{st})}{v_{max}}\right)^{\alpha_Q}} & \text{otherwise} \end{cases}$$

$$k_{p_v} = \kappa \cdot \kappa_0 \cdot q_v \cdot m_0 \cdot G_{star}$$

$$k_{m_v} = \kappa_0 \cdot q_v \cdot (1 - \kappa \cdot m_0 \cdot G_{star})$$

$$K_{\kappa} = k_{p_v} \cdot M_A \cdot n_1 \cdot L_{oz} \cdot (1 - N) - k_{m_v} \cdot N$$

$$\frac{\mathrm{d}N}{\mathrm{d}time} = K_{\kappa}$$

Partial conversion from CellML 1.0 to TEX was done using COR (0.9.31.1409)
Copyright 2002-2018 Dr Alan Garny
